# Absolute number of three populations of interneurons and all GABAergic synapses in the human hippocampus

**DOI:** 10.1101/2023.09.26.559559

**Authors:** Virág Takács, Péter Papp, Áron Orosz, Zsuzsanna Bardóczi, Tamás Zsoldos, Krisztián Zichó, Masahiko Watanabe, Zsófia Maglóczky, Péter Gombás, Tamás F. Freund, Gábor Nyiri

## Abstract

The human hippocampus, essential for learning and memory, is implicated in numerous neurological and psychiatric disorders, each linked to specific neuronal subpopulations. Advancing our understanding of hippocampal function requires computational models grounded in precise quantitative neuronal data. While extensive data exist on the neuronal composition and synaptic architecture of the rodent hippocampus, analogous quantitative data for the human hippocampus remain very limited. Given the critical role of local GABAergic interneurons in modulating hippocampal functions, we employed unbiased stereological techniques to estimate the density and total number of three major GABAergic cell types in the human hippocampus: parvalbumin (PV)-expressing, somatostatin (SOM)-positive, and calretinin (CR)-positive interneurons. Our findings reveal an estimated 49,400 PV-positive, 141,500 SOM-positive, and 250,600 CR-positive interneurons per hippocampal hemisphere. Notably, the higher proportion of CR-positive, primarily interneuron-selective, cells in humans, compared to rodents, may enhance local interneuron regulation. Additionally, using 3-dimensional electron microscopy, we estimated approximately 25 billion GABAergic synapses per hippocampal hemisphere, with PV-positive boutons comprising around 3.5 billion synapses, or 14% of the total GABAergic synapses. These findings contribute crucial quantitative insights for modeling human hippocampal circuits and understanding its complex regulatory dynamics.

## INTRODUCTION

Characterizing the structure, neuronal connections and ultimately the operating principles of the human hippocampus is essential for understanding its involvement in the formation, consolidation, and retrieval of memory traces, in supporting spatial navigation, in adaptive behaviors, and in various cognitive functions (Cai et al., 2016; Roy et al., 2017; Szőnyi et al., 2019; Goode et al., 2020; Zichó et al., 2023). These require computational models that are based on accurate data on the number of neurons in different neuronal classes, which are the cornerstone of any type of modeling (Brunton and Beyeler, 2019; Gandolfi et al., 2023).

Unfortunately, there are only a limited number of studies that have attempted to collect statistically reliable data on key quantitative parameters of the human hippocampus (West and Gundersen, 1990; Harding et al., 1998), and in particular, key GABAergic interneurons have hardly been studied. Recently, Gandolfi et al. proposed a full-scale scaffold model of the human hippocampal CA1 area (Gandolfi et al., 2023). The total number of cells within this volume was estimated to be 18 million, including glial cells and neurons. Based on the assumption that the cellular composition of human and rodent hippocampi is similar, they calculated that 5.28 million of these cells would be neurons, out of which 4.8 million would be principal cells and 0.48 million would be GABAergic interneurons. However, no direct estimate of the different subpopulations of interneurons has been published.

Different inhibitory interneuron subtypes in the human and rodent hippocampi project to different layers, providing functional segregation of inhibitory patterns (Freund and Buzsáki, 1996; Klausberger and Somogyi, 2008; Pelkey et al., 2017). Inhibitory cells of diverse origin and function exhibit distinct protein expression patterns, allowing the classification of hippocampal interneurons into subgroups based on specific molecular markers (Freund and Buzsáki, 1996; Klausberger and Somogyi, 2008; DeFelipe et al., 2013; Pelkey et al., 2017). A large subset of interneurons express the calcium-binding molecule parvalbumin (PV), and most of these cells preferentially innervate the perisomatic regions of principal cells: the cell body and the proximal dendrites [basket cells, (Katsumaru et al., 1988)] or the axon initial segments [axo-axonic cells; (Somogyi, 1977; Somogyi et al., 1982; DeFelipe et al., 1989)]. Expression of the neuropeptide somatostatin (SOM) can be detected in a significant proportion of dendrite-targeting interneurons (Kosaka et al., 1988; Banovac et al., 2022), which preferentially target distal dendrites of principal cells (Klausberger, 2009; Honoré et al., 2021; Takács et al., 2024). A large population of interneurons produce another calcium-binding protein, calretinin (Barinka and Druga, 2010). Many of these neurons are interneuron-selective interneurons that innervate other interneurons and thus can directly regulate inhibition within the hippocampus (Gulyás et al., 1996; Urbán et al., 2002; Tóth et al., 2010). These interneurons are ideally positioned to orchestrate the formation and reactivation of memory traces, to control the flow of information, and to establish and maintain hippocampal rhythms (Klausberger and Somogyi, 2008; Josselyn and Frankland, 2018; Pignatelli et al., 2019). Therefore, basic statistical data on the human hippocampal interneurons are necessary and are highly valuable for building models of this network.

This is particularly important for a better understanding of neurological and psychiatric disorders associated with different subpopulations of hippocampal neurons in Alzheimer’s disease, major depressive disorder, schizophrenia or temporal lobe epilepsy. In certain neurological and psychiatric disorders, changes in the number, distribution, or connectivity of specific interneuronal subgroups expressing different markers in the hippocampus have been described (Maglóczky et al., 2000; Pelkey et al., 2017; Ruden et al., 2021; Vid Prkačin et al., 2023). For example, the density of PV-positive hippocampal interneurons is reduced in patients with schizophrenia (Zhang and Reynolds, 2002), which may play an important role in the deficits observed in gamma oscillations (Lisman et al., 2008; Marín, 2012). The density of calretinin (CR)-expressing interneurons is reduced in epileptic patients, potentially leading to reduced local control over hippocampal interneurons (Maglóczky et al., 2000; Tóth et al., 2010). According to animal models, malfunctioning of SOM-positive interneurons can also contribute to the network alterations occurring in epilepsy (Cossart et al., 2001; Drexel et al., 2022). Both SOM- and CR-positive interneurons have been shown to be vulnerable to ischemic injury (Johansen et al., 1987; Freund and Maglóczky, 1993), and some interneurons may also be differentially affected in Alzheimer’s disease (Ramos et al., 2006; Palop and Mucke, 2016; Waller et al., 2020; Honoré et al., 2021). Therefore, understanding these pathomechanisms requires determining the number and distribution of interneuron types in the hippocampus of healthy subjects.

Here, we estimated the total number of PV-, SOM- and CR-positive inhibitory interneurons in the human hippocampus. We also estimated the total number of GABAergic terminals and synapses in different layers of the hippocampus and the proportion of synapses formed by PV-positive cells. Since the immunohistochemical detectability of certain proteins deteriorates rapidly with prolonged postmortem time (Wittner et al., 2005; Tóth et al., 2010; Gonzalez-Riano et al., 2017), a major advantage of this investigation was that we used fixative-perfused human brains with a short postmortem delay (≤ 3.5 hours).

## RESULTS

### Volume of subregions and layers in one hippocampal hemisphere

Preparing post-mortem human hippocampal samples that retain high immunogenicity for quality immunohistochemistry is inherently challenging, which often makes examining specific neuronal populations difficult or unfeasible. To ensure reliable sample quality and accurately capture the true variability in interneuronal data with stereological precision, first we tested fixative-perfused human brain samples with minimal post-mortem delay (less than 3.5 hours) from 10 individuals. Recognizing the frequent differences in preservation quality even between hemispheres, we analysed each hemispheres independently and selected four hippocampal hemispheres that had the best tissue quality for our study (SKO24, SKO25, SKO27, and SKO31).

For sampling, we employed a non-traditional yet highly precise approach. While conventional stereological methods typically recommend equidistant, parallel, systematic random sectioning (West and Gundersen, 1990) we found that this approach presented two significant challenges: (i) it often made it difficult, if not impossible, to accurately identify the 15 distinct subregions and sublayers in the intricately folded human hippocampus, and (ii) it complicated the handling of very large single sections. The complex folding of hippocampal layers and subregions could introduce substantial biases in our measurements, potentially outweighing the advantages of the traditional method.

To address these concerns, we adopted a design-based approach, primarily sectioning the hippocampus in parallel, with occasional angle adjustments to follow the hippocampal longitudinal axis (Fig. 1). This orientation enhanced the clarity of typical layer structures, facilitated more accurate subregion identification, and improved ease of tissue handling. Although more time-intensive than traditional methods, this alternative approach allowed us to achieve greater precision in measuring the volumes of hippocampal subregions and accurately assessing interneuronal distributions across these regions.

**Fig. 1.**
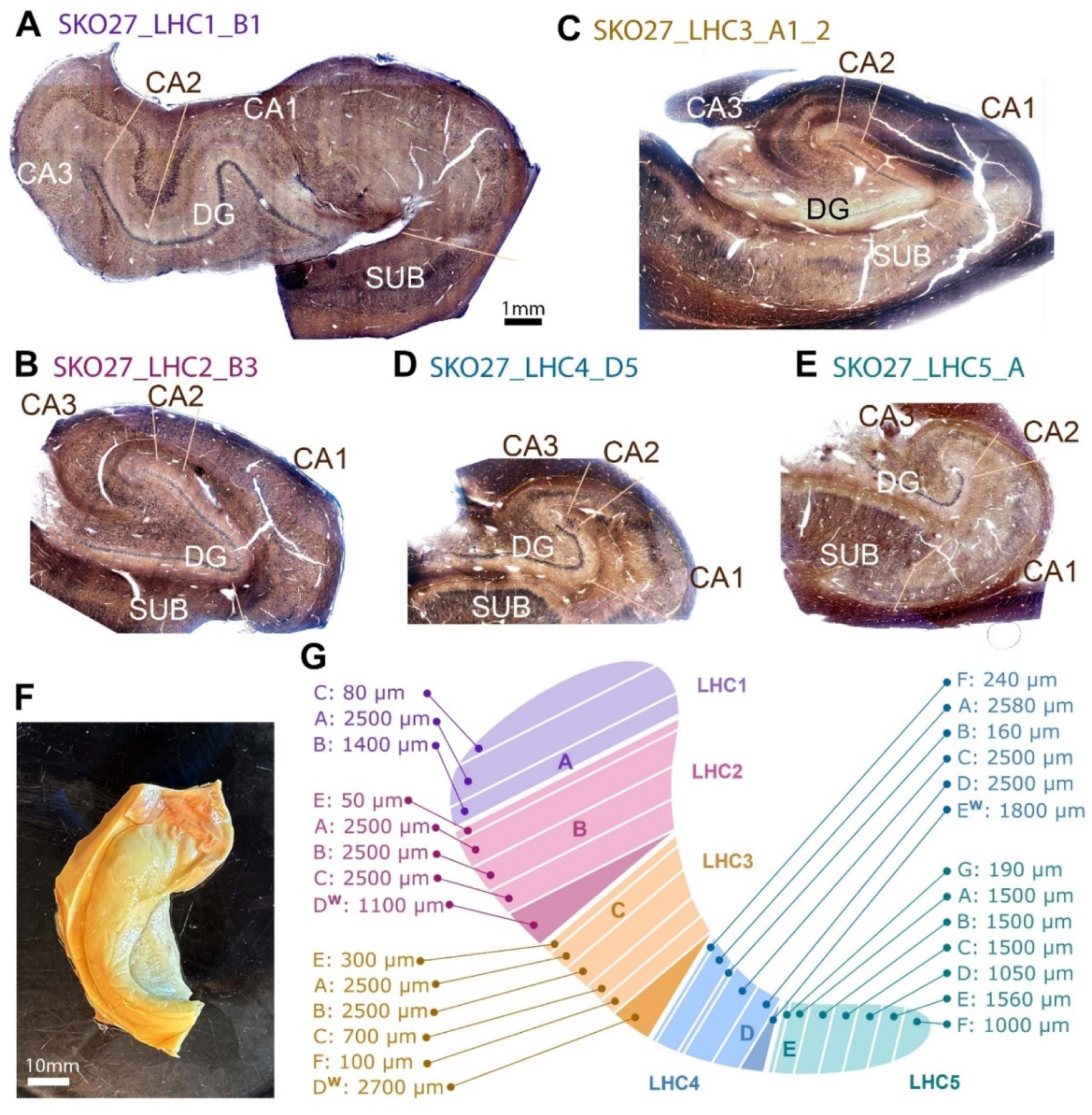
Sectioning procedure. This illustration represents the sectioning procedure (SKO27 is shown here that was similar to the sectioning of the other three hippocampi). Whole hippocampal hemispheres were dissected from the brains (F) and cut into four or five blocks (left hippocampus 1-5 blocks: LHC1-5) perpendicular to their long axis (G). To correctly identify all hippocampal subregions and layers, the curvature of the hippocampus was followed by using 3 precisely reconstructed wedges (LHC2 D^W^, LHC3 D^W,^ and LHC4 E^W^ in G), the volumes of which layers were determined separately (see Methods). All blocks (LHC1-5) were embedded in agarose and completely sectioned while data of section numbers and thickness was accurately recorded for volume measurements (G). A-E: Light microscopic images of 100 µm-thick human hippocampal sections labeled for parvalbumin. The cutting positions of the sections in A-E are indicated in G with letters A-E. Sections from the sampling areas (n=10, for each immunolabeling, including sections in A-E) were used to estimate total interneuron numbers in the hippocampus. Some cracks caused by the flat embedding procedure were also reconstructed and taken into account for both volume and cell density estimation. The scale bar in A applies to B-E.DG: dentate gyrus, LHC1-5: left hippocampus block 1-5, SUB: subiculum.

Accordingly, each hippocampus was sectioned into blocks along its longitudinal axis, allowing for precise volumetric calculations of each hippocampal layer within 15 distinct subregions. These calculations spanned the entire longitudinal extent of the hippocampus for all subjects, derived from Neurolucida tracings of 8-30 PV-labeled sections per tissue block, spaced roughly evenly along the hippocampal axis (refer to Methods, Figs. 1 and 2). Subfield boundaries—including CA1-3 and the dentate gyrus (DG)—were determined based on adjacent calbindin-labeled sections (see Methods, Fig. 2D), following specific structural markers: calbindin-positive mossy fiber bundles delineate the CA3 subfield, while dense calbindin expression in pyramidal cells is unique to CA2 (Sloviter et al., 1991; Merino-Serrais et al., 2020). The boundary between CA1 and the prosubiculum was identified at the point where the pyramidal cell layer nearly occupies the entire layer width between the stratum moleculare and the alveus (Palomero-Gallagher et al., 2020).

**Fig. 2.**
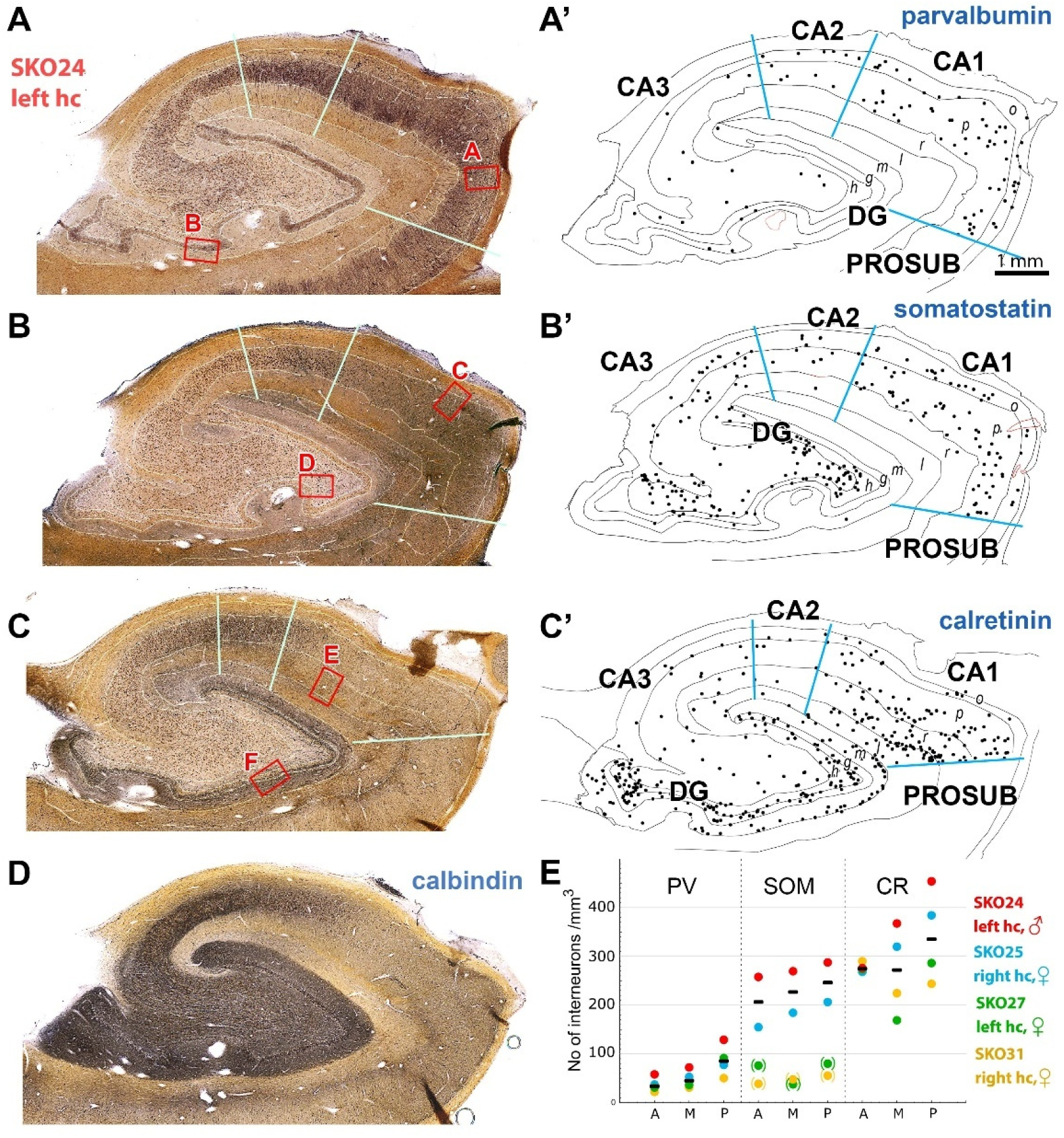
Distribution and cell densities of different types of interneurons in the human hippocampus. Hippocampal sections immunolabeled for parvalbumin (PV) (A), somatostatin (SOM) (B), and calretinin (CR) (C) were used to estimate the total interneuron numbers in the hippocampus. Boundaries of hippocampal subregions in each section were defined using the neighboring calbindin labeled sections (D, see Methods). Four neighboring sections from the left hippocampal hemisphere of SKO24 are shown here to illustrate the characteristic differences in the distribution and density of the different interneurons. The areas highlighted with red are illustrated in Fig. 3 at higher magnification. A’-C’: Neurolucida drawings of hippocampal sections A-C. DABNi-labeled cell bodies were counted using stereological rules (see Methods) and indicated in the drawings with black dots. Subregion boundaries are shown with blue lines. E: Densities (cell number/ mm^3^) of PV, SOM and CR-labeled interneurons in the anterior-, middle-, and posterior third of the hippocampus (A, M and P) in four subjects. Different colors represent data from different subjects (red: SKO24 left-, blue: SKO25 right-, green: SKO27 left hemisphere, yellow: SKO31 right hemisphere). Medians are shown with black lines. In SKO27 and SKO31, the somatostatin labeling was much weaker than in the other two hippocampi, therefore its data are shown in brackets and only data from the other two subjects were used for the median. DG: dentate gyrus, g: stratum (str.) granulosum, h: hilus, hc: hippocampus, l: str. lacunosum-moleculare, m: str. moleculare, o: str. oriens, p.: str. pyramidale, PROSUB: prosubicuum, r: str. radiatum. The scale bar on A’ applies to A-D and B’-C’.

Differences in hippocampal hemisphere sizes became immediately apparent upon initial examination. Indeed, after measurements, we found large differences in the size of the hippocampi of the four subjects: the four hippocampal hemispheres investigated had volumes of 563 (SKO24), 717 (SKO25) 1107 (SKO27) and 1470 (SKO31) mm^3^ (Extended data Tables). Different subregions contributed differently to the total volume of the hippocampus (Extended Data Tables). The CA1 was the largest in all subjects (50.1, 31.4, 40.7 and 46.9%), followed by the CA3 (21, 30.9, 26.5 and 21.9%) and the dentate gyrus (17.9, 26.8, 26.8 and 18.8%) and finally the CA2 (11, 10.9, 6 and 12.4 %), in SKO24, SKO25, SKO27, and SKO31 respectively.

### Estimation of the total number of different interneurons in the human hippocampus

To estimate the total number of specific interneurons, we used whole hippocampal sections immunostained for PV, SOM, or CR using ammonium nickel sulfate-intensified 3-3’-diaminobenzidine (DABNi) (Fig. 1-3, see Methods).

**Fig. 3.**
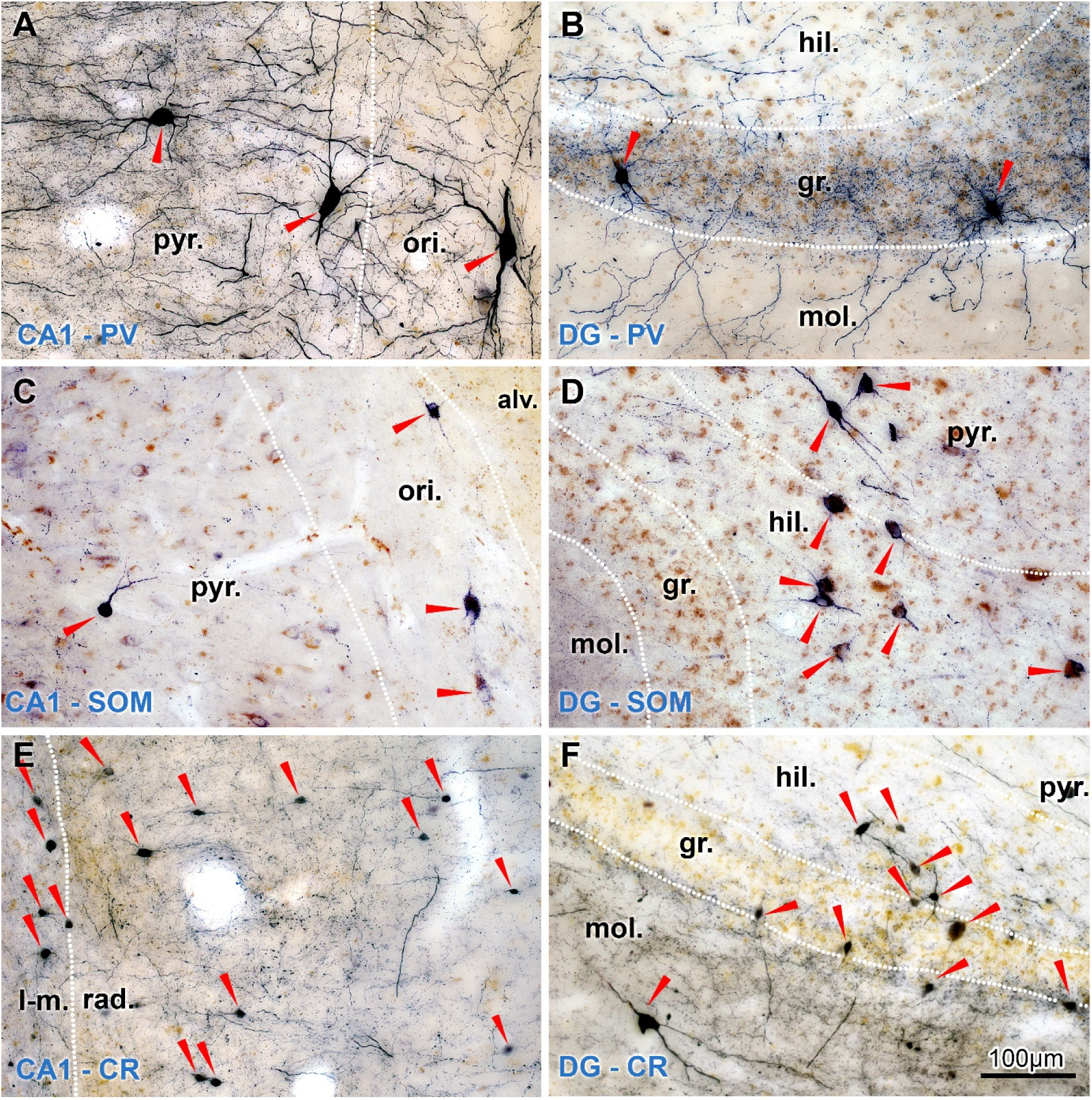
Parvalbumin (PV)-, somatostatin (SOM)- and calretinin (CR)-positive cells in the CA1 area and the dentate gyrus of the human hippocampus. Areas highlighted with red rectangles in Fig. 2 are illustrated here. Cells were immunolabeled using ammonium nickel sulfate-intensified 3-3’-diaminobenzidine (DABNi) resulting in a bluish-black color, which could be easily distinguished from the light brown background color of cell bodies (see e.g. granule cell bodies in str. granulosum). Positive interneurons are indicated with red arrowheads. alv: alveus, DG: dentate gyrus, gr.: stratum (str.). granulosum, hil.: hilus, l-m.: str. lacunosum-moleculare, mol.: str. moleculare ori.: str. oriens; pyr.: str. pyramidale, rad.: str. radiatum. The scale bar in F applies to A-E.

First, we measured the total volume of all 15 layers in each hippocampal section as described above in either 60 or 100 µm thick sections that were evenly distributed along the longitudinal axis of the hippocampus. Then, cell counting was performed on altogether 94 complete sections from all subjects for SOM, PV and CR. Immunolabeled cell bodies were quantified using the Neurolucida software following all stereological principles (see Methods, Fig. 2). Cell density values (number of cells/µm^3^) were calculated for each layer of each hippocampal region (CA1-3, DG, Extended Data Tables). Total cell numbers of each layer were calculated separately for each subject by multiplying the cell densities with the respective volume values. Because the hippocampus have different afferent connections and functions along its longitudinal axis (head-body-tail in human hippocampus or ventral-dorsal hippocampus in rodents), therefore their interneuron composition may also differ (Strange et al., 2014). Therefore, in addition to the data for the entire hippocampus (Fig. 4, Table 1, Extended Data Table 1), we also provided the total density and number of PV-, SOM-, and CR-positive inhibitory cells located in the anterior- (head), middle (body), and posterior (tail) thirds of the hippocampal volume (Fig. 2E, Extended Data Tables 2 and 3). Furthermore, in Extended Data Table 1, we presented the cell density values and total cell numbers separately for four or five consecutive blocks and all the 15 layers along the longitudinal axis of the hippocampus in the head-body-tail direction to represent potential finer differences.

**Fig. 4.**
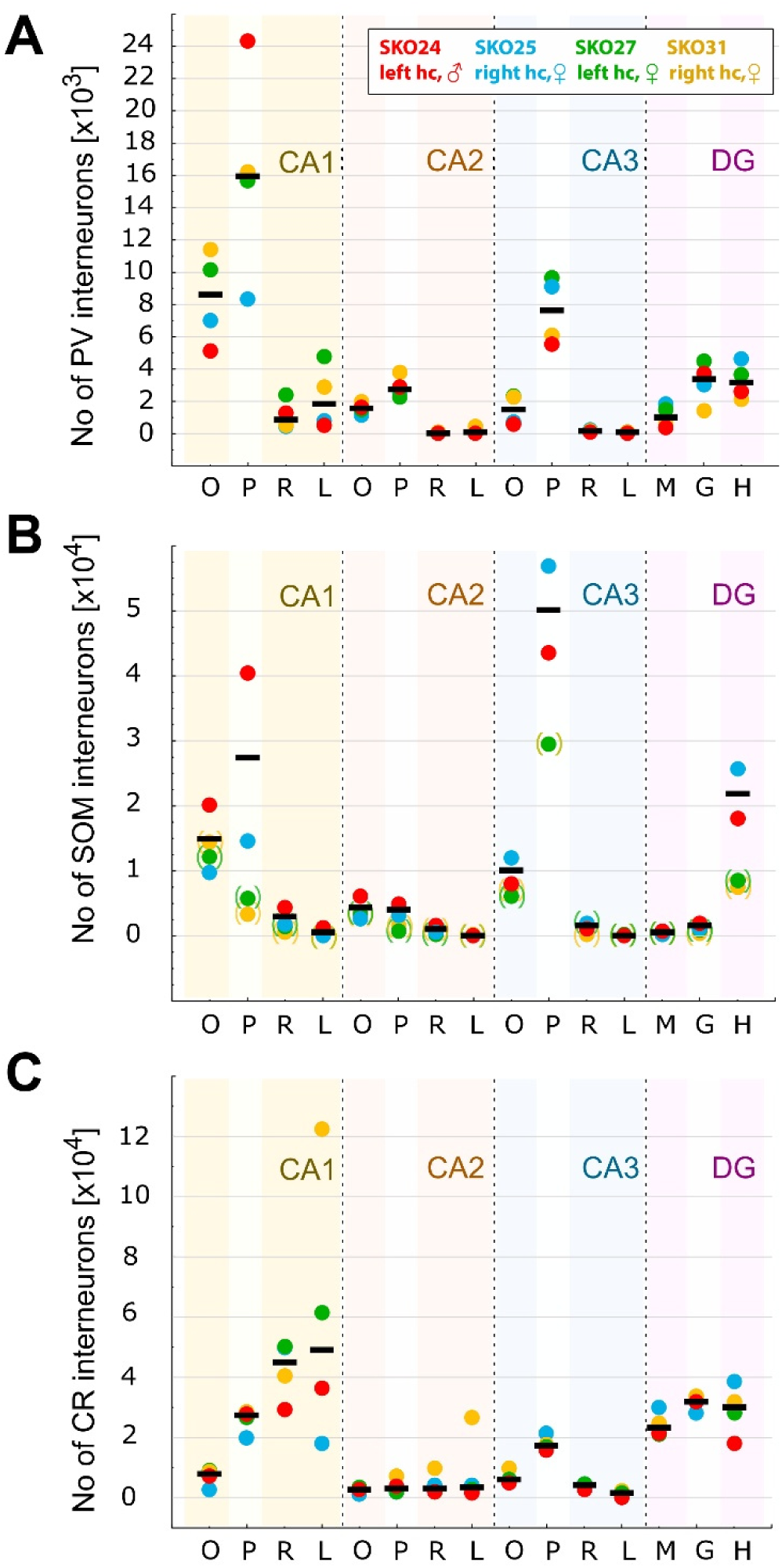
Total numbers of different types of interneurons in the human hippocampus. A-C: Absolute numbers of parvalbumin- (A), somatostatin (B)- and calretinin-positive (C) interneurons in hippocampal layers in one hemisphere. Different colors represent data from different subjects (red: SKO24 left-, blue: SKO25 right-, green: SKO27 left hemisphere, yellow: SKO31 right hemisphere). Medians are shown with black lines. In SKO27 and SKO31, the somatostatin labeling was much weaker than in the other two hippocampi, therefore its data are shown in brackets and only data from the other two subjects were used for the median. DG: dentate gyrus, g: stratum (str.) granulosum, h: hilus, l: str. lacunosum-moleculare, m: str. moleculare o: str. oriens, p: str. pyramidale, r: str. radiatum.

**Table 1.** Estimated total number of PV, SOM, CR interneurons in one human hippocampal hemisphere (PV, CR: medians of four subjects, SOM: medians of two subjects)

| **PV** | **Volume**<br>**[mm<sup>3</sup>]** | **Abs. No** | **SOM** | **Volume**<br>**[mm<sup>3</sup>]** | **Abs. No** | **CR** | **Volume**<br>**[mm<sup>3</sup>]** | **Abs. No** |
| --- | --- | --- | --- | --- | --- | --- | --- | --- |
| CA1 ori | 59.67 | **8611** | CA1 ori | 41.71 | **14963** | CA1 ori | 59.67 | **7923** |
| CA1 pyr | 145.24 | **15955** | CA1 pyr | 99.72 | **27484** | CA1 pyr | 145.24 | **27338** |
| CA1 rad | 79.04 | **893** | CA1 rad | 58.35 | **3063** | CA1 rad | 79.04 | **45050** |
| CA1 lm | 83.58 | **1836** | CA1 lm | 53.78 | **644** | CA1 lm | 83.58 | **48941** |
| CA2 ori | 16.09 | **1567** | CA2 ori | 15.14 | **4361** | CA2 ori | 16.09 | **2570** |
| CA2 pyr | 18.13 | **2733** | CA2 pyr | 22.07 | **4069** | CA2 pyr | 18.13 | **2926** |
| CA2 rad | 19.86 | **0** | CA2 rad | 20.94 | **1068** | CA2 rad | 19.86 | **3274** |
| CA2 lm | 13.18 | **117** | CA2 lm | 11.99 | **43** | CA2 lm | 13.18 | **3483** |
| CA3 ori | 47.94 | **1484** | CA3 ori | 31.20 | **10026** | CA3 ori | 47.94 | **5912** |
| CA3 pyr | 174.69 | **7612** | CA3 pyr | 120.79 | **50168** | CA3 pyr | 174.69 | **17265** |
| CA3 rad | 21.34 | **166** | CA3 rad | 14.06 | **1590** | CA3 rad | 21.34 | **4398** |
| CA3 lm | 8.15 | **93** | CA3 lm | 3.91 | **0** | CA3 lm | 8.15 | **1530** |
| DG mol | 132.00 | **1019** | DG mol | 79.29 | **520** | DG mol | 132.00 | **23153** |
| DG gra | 45.82 | **3379** | DG gra | 33.52 | **1572** | DG gra | 45.82 | **32051** |
| DG hil | 50.71 | **3144** | DG hil | 33.67 | **21957** | DG hil | 50.71 | **29954** |
| Sum | **912** | **49424** | Sum | **640** | **141528** | Sum | **912** | **250576** |

**Table 2.** Estimated total number of vGAT and PV-positive boutons and synapses in the human hippocampus, one hemisphere (data from SKO24)

| Subregion | Layer | Volume (mm<sup>3</sup>) | Labeling | Number of boutons/<br>10<sup>3</sup> μm<sup>3</sup> | Number of synapses/<br>10<sup>3</sup> μm<sup>3</sup> | Abs. No. of boutons<br>*10<sup>6</sup> | Abs. No. of synapses<br>*10<sup>6</sup> |
| --- | --- | --- | --- | --- | --- | --- | --- |
| CA1 | ori | 44.3 | vGAT | 8.55 | 9.26 | 378.3 | 409.8 |
|  |  |  | PV | 0.22 | 0.24 | 9.8 | 10.6 |
|  | pyr | 112.7 | vGAT | 37.58 | 45.09 | 4233.3 | 5079.9 |
|  |  |  | PV | 16.44 | 19.72 | 1851.5 | 2221.8 |
|  | rad | 56.7 | vGAT | 24.00 | 26.00 | 1359.9 | 1473.2 |
|  |  |  | PV | 0.19 | 0.20 | 10.7 | 11.6 |
|  | l-m | 68.3 | vGAT | 37.36 | 40.47 | 2549.9 | 2762.4 |
|  |  |  | PV | 0.02 | 0.03 | 1.6 | 1.8 |
| CA2 | ori | 14.7 | vGAT | 26.36 | 28.56 | 386.5 | 418.7 |
|  |  |  | PV | 0.07 | 0.07 | 1.0 | 1.1 |
|  | pyr | 24.3 | vGAT | 40.29 | 48.34 | 978.3 | 1174.0 |
|  |  |  | PV | 5.37 | 6.44 | 130.4 | 156.4 |
|  | rad | 13.8 | vGAT | 38.16 | 41.34 | 524.7 | 568.5 |
|  |  |  | PV | 0.19 | 0.20 | 2.6 | 2.8 |
|  | l-m | 9.3 | vGAT | 49.52 | 53.65 | 462.9 | 501.5 |
|  |  |  | PV | 0.04 | 0.04 | 0.3 | 0.4 |
| CA3 | ori | 22.3 | vGAT | 22.39 | 24.25 | 499.3 | 540.9 |
|  |  |  | PV | 0.10 | 0.11 | 2.3 | 2.5 |
|  | pyr | 83.4 | vGAT | 27.57 | 33.09 | 2299.0 | 2758.8 |
|  |  |  | PV | 4.75 | 5.70 | 396.1 | 475.4 |
|  | rad | 9.6 | vGAT | 24.64 | 26.69 | 237.0 | 256.8 |
|  |  |  | PV | 0.10 | 0.11 | 1.0 | 1.0 |
|  | l-m | 3.1 | vGAT | 24.88 | 26.95 | 76.6 | 83.0 |
|  |  |  | PV | 0.12 | 0.13 | 0.4 | 0.4 |
| DG | mol | 52.8 | vGAT | 47.93 | 50.01 | 2528.5 | 2638.4 |
|  |  |  | PV | 0.06 | 0.06 | 2.9 | 3.1 |
|  | gra | 26.3 | vGAT | 28.30 | 37.73 | 744.8 | 993.0 |
|  |  |  | PV | 8.62 | 11.50 | 227.0 | 302.7 |
|  | hil | 21.6 | vGAT | 21.47 | 22.40 | 462.8 | 482.9 |
|  |  |  | PV | 0.20 | 0.21 | 4.4 | 4.6 |
| SUM |  | 563 | vGAT |  |  | 17722 | 20142 |
|  |  |  | PV |  |  | 2642 | 3196 |

**Table 3.** Estimated total number of vGAT and PV-positive boutons and synapses in the human hippocampus, one hemisphere (data from SKO25)

| Subregion | Layer | Volume (mm<sup>3</sup>) | Labeling | Number of boutons/<br>10<sup>3</sup> μm<sup>3</sup> | Number of synapses/<br>10<sup>3</sup> μm<sup>3</sup> | Abs. No. of boutons<br>*10<sup>6</sup> | Abs. No. of synapses<br>*10<sup>6</sup> |
| --- | --- | --- | --- | --- | --- | --- | --- |
| CA1 | ori | 39.2 | vGAT | 26.02 | 28.19 | 1019.1 | 1104.0 |
|  |  |  | PV | 0.47 | 0.50 | 18.2 | 19.7 |
|  | pyr | 86.8 | vGAT | 48.78 | 58.54 | 4233.7 | 5080.5 |
|  |  |  | PV | 16.36 | 19.63 | 1419.8 | 1703.7 |
|  | rad | 60.0 | vGAT | 41.84 | 45.32 | 2511.6 | 2720.9 |
|  |  |  | PV | 0.16 | 0.18 | 9.9 | 10.7 |
|  | l-m | 39.3 | vGAT | 46.19 | 50.04 | 1815.8 | 1967.1 |
|  |  |  | PV | 0.15 | 0.17 | 6.0 | 6.5 |
| CA2 | ori | 15.6 | vGAT | 12.72 | 13.78 | 198.7 | 215.3 |
|  |  |  | PV | 0.31 | 0.33 | 4.8 | 5.2 |
|  | pyr | 19.9 | vGAT | 24.64 | 29.56 | 489.1 | 586.9 |
|  |  |  | PV | 14.51 | 17.41 | 287.9 | 345.5 |
|  | rad | 28.1 | vGAT | 23.08 | 25.01 | 649.3 | 703.4 |
|  |  |  | PV | 0.21 | 0.22 | 5.8 | 6.3 |
|  | l-m | 14.6 | vGAT | 59.51 | 64.47 | 870.6 | 943.1 |
|  |  |  | PV | 0.17 | 0.19 | 2.5 | 2.7 |
| CA3 | ori | 40.1 | vGAT | 17.94 | 19.43 | 719.2 | 779.1 |
|  |  |  | PV | 0.09 | 0.10 | 3.7 | 4.1 |
|  | pyr | 158.2 | vGAT | 35.84 | 43.01 | 5669.6 | 6803.5 |
|  |  |  | PV | 6.60 | 7.92 | 1043.8 | 1252.6 |
|  | rad | 18.5 | vGAT | 46.57 | 50.45 | 861.8 | 933.6 |
|  |  |  | PV | 0.16 | 0.17 | 2.9 | 3.2 |
|  | l-m | 4.7 | vGAT | 52.62 | 57.00 | 249.1 | 269.8 |
|  |  |  | PV | 0.05 | 0.06 | 0.3 | 0.3 |
| DG | mol | 105.8 | vGAT | 50.67 | 52.87 | 5361.7 | 5594.8 |
|  |  |  | PV | 0.14 | 0.14 | 14.3 | 14.9 |
|  | gra | 40.7 | vGAT | 26.36 | 35.15 | 1073.2 | 1430.9 |
|  |  |  | PV | 6.61 | 8.81 | 269.0 | 358.6 |
|  | hil | 45.8 | vGAT | 14.00 | 14.61 | 641.0 | 668.8 |
|  |  |  | PV | 0.37 | 0.38 | 16.8 | 17.5 |
| SUM |  | 717 | vGAT |  |  | 26363 | 29802 |
|  |  |  | PV |  |  | 3106 | 3752 |

### The total number of parvalbumin-positive interneurons in the human hippocampus

The distribution and morphology of PV-positive cells were similar in the four subjects (Fig.2 A, A’, E, Fig. 3. A, B) and were consistent with some partial data reported earlier (Braak et al., 1991; Sloviter et al., 1991; Seress et al., 1993a; Wittner et al., 2001; Andrioli et al., 2007). Most of the PV-positive cells exhibited intense labeling and a large size with elongated dendrites (Fig. 3 A, B). Among the three cell groups investigated, the number of PV-positive cells was the lowest in all subregions of the hippocampus in all subjects. The absolute number of PV-positive cells was 49,424 cells/ hemisphere, (median, Figs. 2E, 4A, Table 1) significantly smaller than the number CR-positive cells, see below (p=0.030, Mann-Whitney U test). In the CA regions, the majority (95, 94, 85 and 91% in SKO24, 25, 27 and 31, respectively) of the cell bodies were in the str. pyramidale and oriens (Fig. 2 A’, E). PV-positive cells were primarily found in the granule cell layer and in the hilus within the dentate gyrus (95, 81, 85 and 86% in SKO24, 25, 27 and 31, respectively; Fig. 4A, Table 1, Extended Data Table 1).

### The total number of somatostatin-positive interneurons in the human hippocampus

Immunostaining for SOM neurons presented significant challenges, likely due to the lower expression levels of SOM, a neuropeptide, compared to PV and CR, both of which are calcium-binding proteins. Additionally, SOM tends to be stored in large vesicles, preferentially transported to specific neuronal compartments, further complicating detection. Despite these hurdles, after carefully optimizing the labeling protocols, reliable SOM labeling was achieved in two hippocampal samples. In the SOM-labeled sections, DABNi-positive cells were faintly labeled or appeared as dotted, with visible labeling primarily limited to the somata and proximal dendrites (see Fig. 3 C, F).

Upon analysis, however, SOM labeling remained suboptimal for samples SKO27 and SKO31 (refer to Fig. 2 E and 4 B), where labeled cells exhibited weaker signals compared to those in the other two samples. Consequently, while data for SKO27 and SKO31 are reported for completeness, they were excluded from the final median calculation of the total SOM-positive cell count, as indicated in brackets in Fig. 2 F.

The distribution of SOM-positive cells (Fig. 2 B, B’, E, 4 B) was consistent with some partial data reported in the literature (Chan-Palay, 1987). Briefly, the highest cell density (number of cells/µm^3^, Extended Data tables) was observed in the hilus and in CA3 str. pyramidale close to the hilus (40 and 63% of all hippocampal SOM cells in SKO24 and 25, respectively; Extended Data Tables, Fig 2 B, B’, F, Fig. 3. C). Relatively high numbers of immunolabeled cells were found in str. oriens of CA1-3 (22 and 19% of all hippocampal SOM cells in SKO24 and 25, respectively) and in the CA1 str. pyramidale (26 and 11% of all hippocampal SOM cells in SKO24 and 25, respectively; Fig. 3D). Like PV-positive cells, almost no SOM-positive cells were found in the str. lacunosum-moleculare and radiatum in the CA region and in the str. moleculare of the dentate gyrus. The total number of SOM-positive cells in one hippocampal hemisphere was 2.9 times greater than that of parvalbumin-positive cells (n=141528, median, Figs 2E, 4B, Table 1, Extended Data Table 1).

### The total number of calretinin-positive interneurons in the human hippocampus

CR-positive cells are sensitive to ischaemic conditions (Freund and Maglóczky, 1993), therefore their number rapidly decreases with the post-mortem delay (Tóth et al., 2010). In this study, we utilized samples with minimal post-mortem delay, which contributed to the consistently high quality of immunolabeling across the four subjects, where most cells exhibited prominently labeled cell bodies and dendritic arbors (see Fig. 3 E, F). The location and anatomical characteristics of immunolabeled cells aligned closely with those previously documented (Seress et al., 1993b; Nitsch and Ohm, 1995). Individual CR-positive cells demonstrated considerable morphological diversity, with a majority being relatively small, although some larger cells were also observed (Fig. 3 E, F).

A significant number of CR-positive cells were identified across all layers of the dentate gyrus (see Figs 2 C’, 4C, Table 1, Extended Data Tables). The stratum moleculare of the dentate gyrus was particularly rich in CR-positive cells but contained few, if any, PV or SOM interneurons. CR-positive cells were present in all layers of the CA regions and proved to be the most prevalent of the three interneuron types examined. Their numbers significantly exceeded those of PV-interneurons (p=0.030, Mann-Whitney U test), their median being approximately 5.1 times more numerous than PV-positive neurons (median count: 250,576; see Fig. 2E, Table 1, Extended Data Table 1).

### The total number of GABAergic boutons in the human hippocampus

To model the network architecture of the human hippocampus, it is important to know the number of GABAergic boutons and their approximate number of synapses in different layers and subregions. We measured these data for all GABAergic boutons and PV-positive axon terminals. Estimation of these number for SOM terminals is not possible because SOM is present only within a few dense core vesicles, resulting in staining only in specific regions of these boutons, where these vesicles are located. In addition, there was a significant variability in the quality of staining and the quantity of stained boutons in different subjects and even in different sections of the same subjects. In addition, quantification of CR-containing axonal terminals would not have been particularly useful for modelling purposes because there are a large number of excitatory (and possibly inhibitory) afferent CR-axons that originate from external sources (Amaral and Cowan, 1980; Maglóczky et al., 1994; Borhegyi and Leranth, 1997; Bokor et al., 2002; Urbán et al., 2002; Tóth et al., 2010).

### The total number of vGAT-positive boutons and synapses in the human hippocampus

To quantify the total number of GABAergic boutons, we employed immunoperoxidase labeling for the vesicular GABA transporter (vGAT; see Methods). For this analysis, we selected sections from subjects SKO24 and SKO25, as they demonstrated the highest vGAT labeling density alongside acceptable ultrastructure preservation for quantitative analysis. In these sections, DABNi-labeled, vGAT-positive boutons were manually outlined and counted in three dimensions, following stereological guidelines, using stacks of serial electron microscopy images (refer to Methods, Fig. 5 A, A’).

**Fig. 5.**
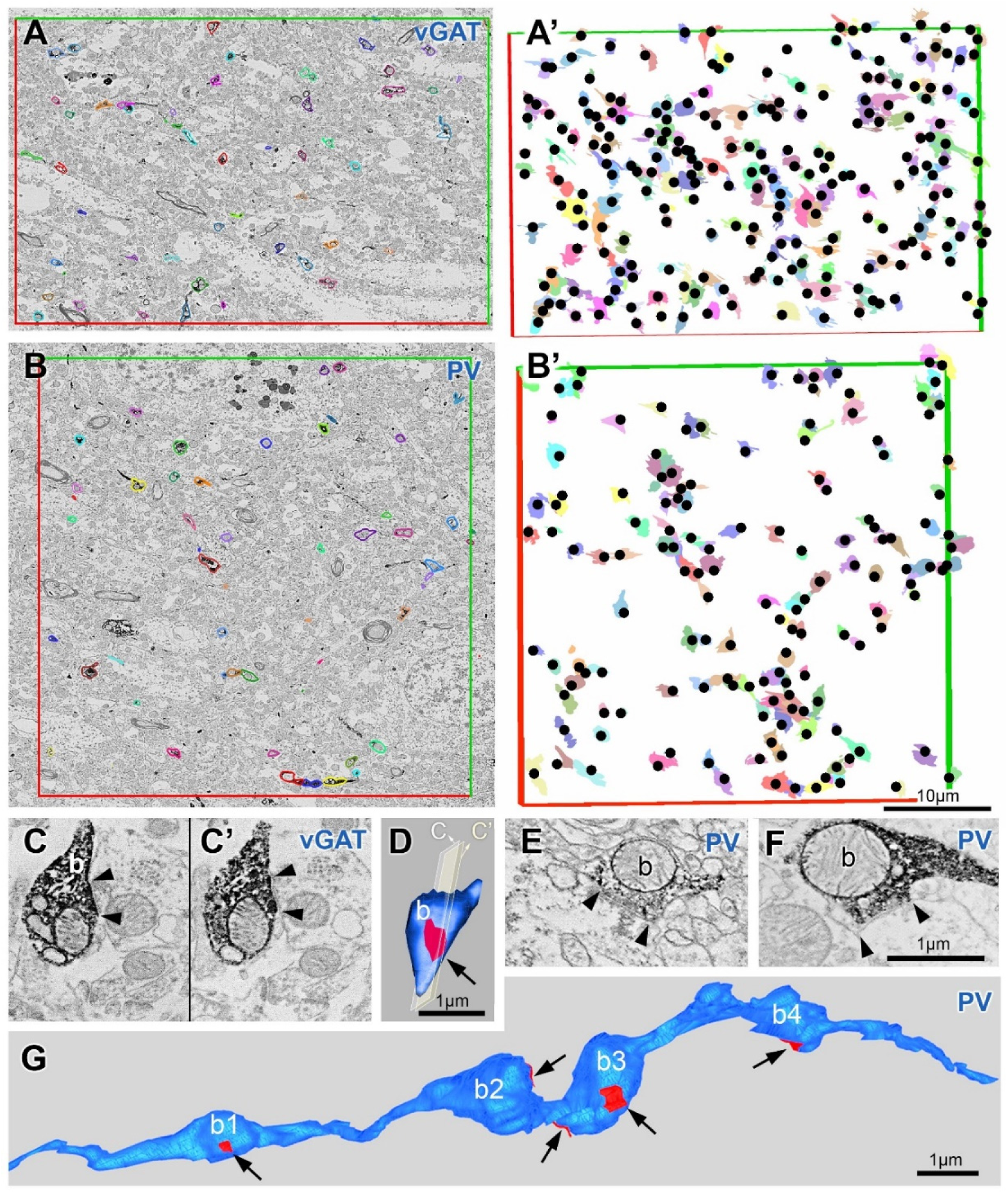
Electron microscopic analysis of vesicular GABA-transporter (vGAT) and parvalbumin (PV) -positive boutons in the human hippocampus. A, B: Electron micrographs of randomly selected areas from CA1 str. pyramidale immunolabeled for vGAT (A) or PV (B). These areas were imaged in every section of a series of 70 nm-thick ultrathin sections (n= 53 serial sections) using a scanning electron microscope. vGAT or PV-positive boutons were manually labeled throughout the series and quantified in three dimensions using stereological rules (red-green rectangle counting frame: exclusion and inclusion lines, respectively). A’ and B’: Three-dimensional view of all vGAT (A’)- and PV-positive (B’) boutons in the sampled volume. Every labeled bouton is indicated with a colored contour and a black dot in A’ and B’. The bouton density of hippocampal layers was multiplied by the volume of the given layer for an estimation of the total number of vGAT or PV-positive boutons in the hippocampus. C-G: Estimation of the number of synapses/boutons. vGAT or PV-positive boutons were reconstructed in 3 dimensions from serial electron microscopic images (see Methods, D, G). Synapses are shown in red and indicated with arrows in the three-dimensional reconstructions. C, E, F: Synapses were identified by a parallel apposition of the pre-and postsynaptic membranes, a widening of the extracellular space at the presumptive synaptic cleft, and a postsynaptic membrane thickening. Arrowheads label the edges of synapses. In the human hippocampus, most of the vGAT and PV-positive boutons formed only one synapse (D, b1, b3, b4 in G) and only occasionally made two synapses (b3 in G). The scale bar in B’ applies to A, A’ and B, and scale bar in F applies to C, C’ and E as well.

The density of GABAergic boutons (expressed as number/mm³) was calculated across sampling volumes ranging from approximately 1,680 to 18,400 µm³ within different hippocampal layers and samples (Tables 2 and 3). Our findings indicate notable variability in bouton density across hippocampal layers and subregions. The stratum lacunosum-moleculare of CA2 displayed the highest density, with 5.95 × 10⁷ boutons/mm³, while the stratum oriens of CA1 exhibited the lowest density, at 8.55 × 10⁶ boutons/mm³.

Given that a single bouton can form multiple synapses (Bourne and Harris, 2012; Takács et al., 2015; Rigby et al., 2023), we conducted a 3-dimensional reconstruction of vGAT-positive boutons (n=36 in dendritic layers of CA1 and n=23 in the stratum moleculare of the dentate gyrus) along with their synapses. This was achieved using electron microscopy image stacks at a resolution optimized for synaptic membrane detection (3 nm/pixel, see Methods, Fig. 4 C-D). Our analysis revealed that most boutons in the human hippocampus formed a single synapse, with only a few exhibiting two synapses. The average number of synapses per bouton was 1.08 in the cornu ammonis and 1.04 in the molecular layer of the dentate gyrus.

To estimate the total number of GABAergic synapses within each hippocampal layer, we multiplied the average synapse number per bouton by the bouton density and the volume of the respective layer in one hemisphere of the same subject (see Tables 2 and 3). It is important to note that due to the limitations of immunocytochemistry, not all vGAT-positive boutons are likely labeled, and ultrastructural preservation in postmortem samples may not allow for the detection of every synapse. Therefore, the values provided in the tables should be regarded as minimum estimates.

### The total number of parvalbumin-positive boutons and synapses in the human hippocampus

Among the parvalbumin-labeled sections, SKO24 and SKO25 displayed the densest axonal labeling, making them ideal for estimating PV-positive bouton density across hippocampal layers. Due to the high density of PV labeling in the somatic layers (see Fig. 3 A, B), we analyzed these layers using electron microscopy stacks, similar to the approach for vGAT-labeled sections (Fig. 4 B, B’). However, in the dendritic layers, PV-positive fibers were sparse, making electron microscopy impractical; instead, we analyzed these layers using light microscopy image stacks created with the Neurolucida system at 100x magnification with an oil immersion objective.

The PV-positive bouton density (expressed as number/µm³) was highest in CA1 stratum pyramidale (1.64 × 10⁷ in both subjects), followed by other perisomatic layers (CA2-3 stratum pyramidale and stratum granulosum; see Tables 2 and 3). Dendritic layers exhibited markedly lower densities (Tables 2 and 3). We calculated the number of synapses per bouton for PV-positive boutons similarly to vGAT-positive boutons, finding that PV-positive boutons in the somatic layers had a higher synapse count per bouton than vGAT-positive boutons in dendritic layers. Specifically, stratum pyramidale exhibited 1.20 synapses per bouton (n=20), while stratum granulosum showed 1.33 synapses per bouton (n=21).

Using these measurements, we calculated the number of synapses formed by PV-positive boutons in each layer and subfield of the hippocampus for both subjects (Tables 3 and 4). Due to the immunocytochemical limitations inherent in detecting PV within axonal boutons in postmortem human tissue, these values represent minimum estimates. Our analysis reveals that PV-positive synapses contribute significantly to the total GABAergic synapse population within the somatic layers of the human hippocampus. Specifically synapses of PV-positive boutons constituted a substantial proportion of all GABAergic synapses in the somatic layers (CA1 stratum pyramidale: 44% and 34%; CA2 stratum pyramidale: 13% and 59%; CA3 stratum pyramidale: 17% and 18%; stratum granulosum: 30% and 25% for SKO24 and SKO25, respectively).

## DISCUSSION

Using unbiased stereological methods, we estimated the absolute numbers of three major interneuron populations, the PV-positive cells, the SOM-positive cells, and the CR-positive cells, in each of the 15 layers of different hippocampal subregions in one hippocampal hemisphere of four human subjects. We found that there are approximately at least 49,400 PV-positive, 141,500 SOM-positive, and 250,600 CR-positive interneurons (median of four subjects for PV- and CR-positive interneurons, median of two subjects for SOM-positive interneurons) in one hemisphere of the human hippocampus (including the cornu ammonis and the dentate gyrus). In addition, we estimated the total number of GABAergic synapses in each layer of all hippocampal subregions using electron microscopy, and we also determined how many of these synapses are formed by parvalbumin-positive cells. We estimate that there are at least 2.5* 10^10^ GABAergic synapses in the entire hippocampus, of which about 3.5* 10^9^ are PV-positive, representing about 14% of all GABAergic synapses (median of two subjects).

### The size of the hippocampus does not correlate with the number of its interneurons

Consistent with previous findings (Harding et al., 1998; Adler et al., 2018; Palomero-Gallagher et al., 2020), we observed substantial variation in hippocampal volumes across the four subjects, ranging from 563 to 1470 mm³, with the largest hippocampus measuring 2.61 times the size of the smallest. Despite this, we found only a 1.8-fold difference in the number of CR-positive interneurons and an almost identical number of PV cells (1.03x difference) between the largest and smallest hippocampi. Notably, while SKO25’s hippocampus was 1.27 times larger than that of SKO24, it contained 17% fewer PV cells and 15% fewer SOM cells.

Given that all sections underwent identical preparation procedures and had a comparably short postmortem delay, these size differences are unlikely to be attributable to varying degrees of tissue shrinkage and likely reflect individual anatomical heterogeneity. Our findings suggest that interneuron numbers are more tightly regulated and less influenced by hippocampal volume variations, indicating that cell numbers may be independently regulated variables. This may reflect differences in glial cell populations, neuronal connectivity, or the presence of other neuronal subtypes, underscoring a potential functional stability in interneuron distribution across individuals.

### Higher proportion of calretinin-containing interneurons in the human hippocampus compared to other interneurons

The three interneuron populations investigated in this study cover only a fraction of all hippocampal interneurons, because there are several other types that do not express any of the markers tested (e.g. calbindin-positive cells and cholecytokinin/ cannabinoid receptor 1 - positive interneurons (Lotstra and Vanderhaeghen, 1987; Seress et al., 1993a; Ludányi et al., 2008)).

We found that the ratio of the three investigated cell groups in the human hippocampus was 1:3:5 (PV:SOM:CR), whereas in rodents this ratio is 2.6:1:1.5 (Bezaire and Soltesz, 2013). While in rodents the parvalbumin-positive cells are the most numerous group, in the human hippocampus the calretinin-positive cells are five times more numerous. Single-cell RNA sequencing analysis has also revealed an evolutionary change in the proportion of inhibitory neuron classes with an increase of caudal ganglionic eminence-derived interneurons (including calretinin-positive interneurons) in the human neocortex (Hodge et al., 2019), confirming previous findings on progenitor cell migration (Hansen et al., 2013). In the primate neocortex, a threefold increase in the proportion of calretinin-positive neurons has also been demonstrated compared to rodents (Džaja et al., 2014; Hladnik et al., 2014).

The abundance of calretinin (CR)-positive interneurons in the human hippocampus is particularly noteworthy due to their distinct functional role among interneurons. In rodents, most CR-positive interneurons are specialized to target other GABAergic cells, acting as modulators within inhibitory networks (Gulyás et al., 1996). Evidence indicates a similar role for human hippocampal CR-positive interneurons, with 39.2% of CR-positive boutons synapsing onto other interneurons, 28% onto unidentified targets, and 25.2% onto pyramidal cells (Urbán et al., 2002). However, the human hippocampus also harbors unique CR-positive cell types that are absent in rodents and have unknown targets, such as multipolar cells at the stratum lacunosum-moleculare border in CA1 and larger neurons in the hilus (Urbán et al., 2002; Tóth et al., 2010). This suggests that the marked increase in CR-positive neurons in humans — paralleling findings in the neocortex (Džaja et al., 2014) — is not merely an expansion of rodent cell types but rather an indicator of evolutionarily novel cells.

Supporting this notion, gene set enrichment analyses comparing human and mouse inhibitory neuron clusters reveal that the human PVALB (parvalbumin) and SST (somatostatin) neuron clusters show substantial gene expression similarities to their mouse counterparts (Ayhan et al., 2021). In contrast, the human VIP/CALB2 (i.e., CR cells) cluster is less aligned, suggesting a more divergent gene expression profile that lacks a direct rodent equivalent (Ayhan et al., 2021). This divergence underscores the need for further investigation into these unique human CR-positive interneurons. Single-cell mRNA sequencing of anatomically identified hippocampal interneurons in human samples could clarify the transcriptomic identity of these evolutionarily new cell types, providing comparative insights with both human and rodent databases (Habib et al., 2017; Ayhan et al., 2021; Tran et al., 2021; Langlieb et al., 2023; Siletti et al., 2023; Zhang et al., 2023).

The increased population of CR-positive interneurons in humans also aligns with recent findings from connectomic studies using 3D electron microscopy, which demonstrate a substantial enhancement in interneuron-to-interneuron connectivity, nearly eight-fold, from rodents to humans (Loomba et al., 2022). This suggests that CR-positive interneurons play a crucial role in forming a more complex and specialized inhibitory network within the human hippocampus, reflecting evolutionary adaptations that may support enhanced regulatory functions in hippocampal circuitry.

### Total number of GABAergic synapses and high proportion of parvalbumin-positive synapses in somatic layers

The density of vGAT-positive synapses in CA1, along with the layer-specific distribution patterns—higher density in the stratum pyramidale and stratum lacunosum-moleculare compared to other layers—closely mirrors observations by Montero-Crespo et al. (Montero-Crespo et al., 2020) on the distribution of symmetric synapses in CA1. In our study, we quantified the density of synapses formed by vGAT-positive and PV-positive boutons across each layer within the entire cornu ammonis and dentate gyrus, allowing us to calculate the total number of boutons and synapses for each hippocampal subregion and layer.

Our findings indicate that a substantial portion of synapses in the somatic layers are contributed by parvalbumin-positive cells, comprising 13.3–58.9% of synapses in the cornu ammonis and 25.1–30.5% in the dentate gyrus. These PV-positive synapses may originate from local interneurons as well as from extra-hippocampal afferents. However, if the majority are established by local PV neurons, this would suggest an impressive capacity of individual PV cells to form extensive axonal networks, highlighting the significant role of PV-positive interneurons in regulating hippocampal circuitry through widespread synaptic connections.

## METHODS

### Human samples

Human hippocampus samples were obtained from four subjects who died due to causes not directly involving any brain disease or damage. All procedures were carried out in compliance with the Declaration of Helsinki and approved by the Regional Committee of Science and Research Ethics of the Scientific Council of Health (ETT TUKEB 31443/2011/EKU, renewed: BM/15092-1/2023). The subjects went through an autopsy in the Department of Pathology of St. Borbála Hospital, Tatabánya, Hungary. Informed consent by relatives was obtained for the use of brain tissue and access to medical records for research purposes. Samples from the following subjects were used in the present study: SKO24: 79 years old, male, post-mortem interval (PMI): 2h 30 min, left hemisphere; SKO25: 87 years old, female, PMI: 3h 30 min, right hemisphere; SKO27: 76 years old, female, PMI: 3h 15 min, left hemisphere, SKO31: 81 years old, female, PMI: 3h, right hemisphere. Brains were removed after death and the internal carotid and vertebrate arteries were cannulated. First, physiological saline containing 0.33% heparin was perfused through this system (1.5 l in 30 min), after which perfusion continued with Zamboni fixative solution containing 4% paraformaldehyde, 0.2% picric acid in 0.1 M phosphate buffer (PB, pH = 7.4, 2 h, 4 l). PMI was determined between the time of death and the start of perfusion by the fixative. Our investigations reveal that the tissue fixation protocol used for human brain perfusion achieved high-quality preservation across the entire hippocampus —suitable for light microscopic immunolabeling measurements— in only approximately 40% of cases. In most instances, preservation quality varied significantly even between the two hippocampal hemispheres, preventing consistent measurements in either the left or right hippocampus across all subjects. Collecting and averaging data from less well-preserved samples would reduce accuracy, making it more meaningful to focus only on the optimally preserved specimens. Achieving the necessary tissue quality for electron microscopy posed an even greater challenge, which is why electron microscopic data in this study were limited to two subjects (SKO24 and SKO25) where preservation was optimal.

### Sectioning

As described in the beginning of the results, for sectioning, we employed a non-classical yet highly precise stereologically unbiased method. Although this approach was more time-consuming than traditional methods, this technique allowed us to achieve greater precision in capturing the volumes of hippocampal subregions and the interneuronal distributions across hippocampal subregions. It is because (i) distinct subregions and sublayers could be differentiated better and (ii) it made handling of very large single sections unnecessary.

Therefore, the whole hippocampi were dissected from the brains and cut into four (SKO24) or five (SKO25, 27, 31) blocks perpendicular to their long axis (Fig. 1). To correctly identify all hippocampal subregions and layers, some sections were cut at an angle, followed by mostly parallel sectioning (see Fig 1). Therefore, in SKO25, 27 and 31 each block ended in a so-called ‘wedge part’, the volumes of which layers were determined separately.

Blocks were embedded into 3% agarose and sections were cut on a Leica VT1200S vibratome. From all blocks, at least one but mostly more (2-3) sampling areas (located 1.5-3 mm away from each other) were sectioned at 60 or 100 µm. Sections in these sampling areas were collected in stereological order. All hippocampal blocks were completely sectioned, sections were collected, and the order, thickness and number of sections were accurately recorded. Sections were rinsed in PB, cryoprotected in 30% sucrose in PB overnight, frozen over liquid nitrogen, and stored at -70 C until further processing.

### Pre-embedding immunolabeling

From all sampling areas four sections were immunostained for PV, SOM, CR, and calbindin, respectively. For electron microscopic measurements selected sections from the medial portion of the hippocampus were immunostained for vesicular GABA transporter (vGAT) or for PV. Sections were freeze-thawed twice over liquid nitrogen in 30% sucrose dissolved in PB. Some sections of SKO31 used for SOM immunolabeling were incubated in 100x citrate buffer (pH 6.0) at 80 Celsius degree for 30 minutes for antigen retrieval. Sections were washed in PB (2x10 min) and treated with 1% sodium borohydride in PB for 10 min. After extensive washes in PB (5x5min) endogenous peroxidase-like activity was blocked by 1% hydrogen peroxide in TBS for 10 min. Then sections were transferred into 0.05 M Tris-buffered saline (pH 7.4; TBS) and blocked in 2% bovine serum albumin (Sigma-Aldrich) and 0.1 g/ml lysine and glycine in TBS. Then, sections were incubated in a solution of mouse anti-PV (1:5000, Swant, Cat. No. 235)/ guinea pig anti-SOM (1:1000, Synaptic Systems, Cat. No. 366 004), mouse anti-CR (1:5000, Swant, Cat. No. 6B3) rabbit anti-calbindin (1:10 000, gift from Prof. Kenneth Baimbridge, code: 8702) or goat anti-vGAT [1:1000, gift from Prof. Masahiko Watanabe (Iwakura et al., 2012; Kudo et al., 2012)] primary antibodies diluted in TBS containing 0.05% sodium azide for 2 days. After repeated washes in TBS, the sections were incubated in horse anti-mouse (Cat. No. BA-2000)/ goat anti-guinea pig (BA-7000)/ goat anti-rabbit (BA-1000) or horse anti-goat (BA-9500) biotinylated secondary antibodies overnight (1:500, Vector Laboratories) followed by extensive washes in TBS (3x10 min) and in VECTASTAIN® Elite® avidin–biotinylated horseradish peroxidase complex (Standard) for 1.5 h (1:250 in TBS; Vector Laboratories).

The immunoperoxidase reaction was developed using ammonium nickel sulfate-intensified 3-3’-diaminobenzidine (DABNi) as the chromogen. Antibodies used in this study were tested for specificity. No specific-like staining was observed without primary antibodies. After washes in PB, sections were contrasted with 0.25% (for cell counting in the light microscope) or 0.5% (for electron microscopic investigations) osmium-tetroxide in PB on ice for 15 min. The sections were then dehydrated in ascending alcohol series and in acetonitrile and embedded in Durcupan (ACM; Fluka). During dehydration, the sections for electron microscopic investigations were treated with 1% uranyl acetate in 70% ethanol for 30 min.

### Cell counting in the light microscope

For each of the three interneuron-populations, 5-10 hippocampal sections (7.8 ± 1.9) at approximately equal distances from each other and in precisely known locations along the longitudinal axis of the hippocampus, representing all blocks (see above) have been processed in each subject (altogether n= 94 sections). Hippocampal layers of PV, SOM and CR-labeled sections were outlined using the Neurolucida software attached to a Zeiss Axioscope2 microscope with a 5× objective (MBF Bioscience). Boundaries of hippocampal subregions (CA1, 2, 3, DG) in each section were defined using the neighboring calbindin labeled sections by applying the following rules: calbindin-positive bundles of mossy fibers indicate the extent of the CA3 subregion, whereas pyramidal cells are densely calbindin-positive only in the CA2 region (Sloviter et al., 1991; Merino-Serrais et al., 2020). The pyramidal cell layer expands and occupies almost all space between the alveus and dentate gyrus in the subiculum which does not apply to CA1 (Palomero-Gallagher et al., 2020). We did not include the prosubiculum in the CA1 region. In addition to references mentioned above, the subfields in the head region were defined using data from (González-Arnay et al., 2024). PV, SOM, or CR-positive cell bodies in CA1-3 and the dentate gyrus were marked on the drawings using the Neurolucida software with a 40x oil-immersion objective. In human samples, OsO4-treatment caused brown background-like labeling of cell bodies that is likely caused by the osmification of lipofuscin. This was clearly distinguishable from the specific DABNi-labeling of interneurons, which was black and labeled the dendrites of the cells as well in addition to their somata (see Fig. 3). All cells were considered positive (even those that were faintly labeled or dotty) if they were black and had at least one dendrite. According to stereological principles, cell bodies were only marked on drawings and counted if they did not intersect the bottom of the section but were counted if cut on the surface. In each sampled section, for each layer of each subregion (CA1, 2, 3: str. oriens, pyramidale, radiatum and lacunosum-moleculare, dentate gyrus: hilus, str. granulosum, and moleculare) a density value was calculated by dividing the cell count obtained by the area of the sampled part of the given section. To obtain the total cell count of a block of the human hippocampus, the densities of the two bounding sampled sections (60 or 100 μm) were averaged and multiplied by the volume of the respective layer in the block. Finally, the cell counts estimated in these blocks were summed to give the total hippocampal cell count in every layer (in every subregion) of the hippocampus. Because we utilized the same sections for both total volume- and cell density measurements, we did not apply adjustments for tissue shrinkage/dilatation.

### Measurement of the volume of hippocampal layers

Neurolucida drawings of PV or calbindin-labeled sections (n=9-30 sections/subjects) in precisely known locations along the long axis of the hippocampus, representing all blocks were used for volume measurements (see above). Subregions (CA1-3, DG) in PV-labeled sections were defined also with the help of the neighboring calbindin-labeled sections (see above). The areas of each hippocampal layer in each subregion (CA1, 2, 3: str. oriens, pyramidale, radiatum and lacunosum-moleculare, dentate gyrus: hilus, str. granulosum, and moleculare), were measured using the Adobe Photoshop software. The volume of each hippocampal layer in each subregion was defined for all tissue blocks. In the “wedge” parts, sections (with precisely known locations along the longitudinal axis of the hippocampus) were labeled for calbindin. The area of hippocampal subregions in each of these wedge sections of SKO24, 25 and 27 was defined using the Cavalieri Estimator probe in StereoInvestigator (MBF Bioscience), whereas in SKO31 we used Neurolucida drawings for these measurements. The volumes of each layer of each block were summed to give the total volume of the given layer in the hippocampal hemisphere.

### Sample preparation for scanning electron microscopy (SEM)

The tissue was cut out and mounted on the top of resin blocks with cyanoacrylate glue, then ultrathin (70 nm thick) sections were cut with a diamond knife (Diatome, Biel, Switzerland) mounted on an ultramicrotome (EM UC6, Leica, Wetzlar, Germany). Ribbons of consecutive ultrathin sections (n= 55-200 sections/sample) were transferred to silicon wafers (a special conductive section carrier, Ted Pella).

### SEM imaging

Because of the very dense DABNi-labelling, the number of DABNi-labeled boutons could not be accurately determined at the light microscopic level in the vGAT-labeled sections and in the somatic layers of the PV-labeled sections. Therefore, we used an electron microscope to obtain these values. The sections were imaged with scanning electron microscopy (SEM, FEI Apreo, Eindhoven, The Netherlands) in backscattered mode. We used a FEI Apreo field emission gun (FEG) SEM (Thermo Scientific) and a T1 in-column detector to record the backscattered electrons. The micrographs were acquired with an array tomography plugin in MAPs software (version: 3.17, https://www.thermofisher.com/hu/en/home/electron-microscopy/products/software-em-3d-vis/maps-software.html), which can facilitate the automatic acquisition of SEM images. The following imaging parameters were used: 4 mm working distance at a high vacuum with 0.1 nA beam current, 2 kV high voltage, 3 μs/pixel dwell time and 9 nm pixel size for bouton density measurements and 6 μs/pixel dwell time and 3 nm pixel size for analysis of number of synapses/bouton. The micrographs were 6000x6000 (bouton density measurements, see below) or 8000x8000 pixels (analysis of number of synapses/bouton, see below) and we recorded 45-127 sections with 70 nm thickness.

### Correction for EM section compression during diamond knife sectioning

The force applied by the diamond knife during the cutting of serial EM sections compresses the sections. The correction factor for section compression was calculated by dividing the width of the tissue block face in the resin with the width of EM sections measured perpendicular to the diamond knife blade (Y plane). The measurements were performed using Zeiss Axioplan2 light microscope. The correction factor in the Y plane was 1.27 ± 0.05 (mean ± SD, n= 15 blocks). There was no change in the EM section dimension in the direction parallel (X plane) with the diamond knife edge. The area of sampled ROI was calculated by multiplying the X length and the corrected Y lengths of the ROI. It was multiplied by the number of sections and section thickness for the calculation of the volume of sampled tissue.

### SEM Image analysis

After SEM image series were collected, image post-processing was done in Fiji ImageJ (Schindelin et al., 2012). The SEM stacks were imported into Fiji TrakEM2 plugin15 and finely aligned. The boutons and synapses were segmented using the IMOD (version 4.9, https://bio3d.colorado.edu/imod) package (Kremer et al., 1996) in which the delineation was performed on a Wacom Cintiq 27QHD Creative Pen and display tablet.

### Bouton density measurements

Randomly selected sampling areas were imaged in every section of a series of 70 nm-thick ultrathin sections (n= 50-127 serial sections) using a scanning electron microscope (see above). The micrographs were 6000x6000 pixels with 9 nm pixel size. After fine alignment of the image series (see above), boutons were manually labeled using IMOD software (see above) in the images throughout the series and quantified in 3D using stereological rules. We used the physical disector principle. Counting was done within a stereological counting frame and boutons were only counted if they were not present in the last section. The sampling volume was 1680-19800 µm^3^ / hippocampal layers/ sample. In layers where the density of PV-labeled boutons was very low (i.e., dendritic layers), we counted the boutons at light microscopic level using stacks of images created with the Neurolucida system attached to a Zeiss Axioscope2 microscope with a 100× immersion oil objective (MBF Bioscience). The sampling volume in the light microscope was 598 000-627 000 µm^3^ / hippocampal layers/ sample.

### Determination of synapse number/ bouton

To determine the number of synapses formed by a single bouton, we used only the SKO24 sample, as in certain layers and areas its ultrastructure was much better preserved than that of SKO25. Randomly selected sampling areas were imaged in every section of a series of 70 nm-thick ultrathin sections (n=45-67 serial sections) using a scanning electron microscope at 3 nm/pixel resolution (see above). DABNi-labeled boutons and their synapses were reconstructed only if they were fully included in the series and the preservation of surrounding tissue was sufficient to identify synapses. Parallel appositions between the membranes of the presynaptic bouton and the putative postsynaptic target were regarded as synapses if they displayed a widening of the extracellular space at the presumptive synaptic cleft and a postsynaptic membrane thickening. Synapse number/ boutons measurements were done in vGAT-labeled sections in the case of dendritic layers and PV-labeled sections in the case of somatic layers.

## ACKNOWLEDGEMENTS

We thank E. Szépné Simon, N. Kriczky, and Z. Hajós for helping with the experiments, Q. Nguyen for helping with Neurolucida drawings and G. Goda and K. Iványi for other assistance. We thank the Department of Pathology, Saint Borbála Hospital, Tatabánya, Hungary and the Human Brain Research Laboratory at the Institute of Experimental Medicine (IEM) for providing human brain tissue. We thank the Electron Microscopy Center at the IEM for technical help with the FEI Apreo scanning electron microscope and the Hitachi H-7100 transmission electron microscope and S. Kőszegi for help in image acquisition. We acknowledge the Light Microscopy Center at the IEM for the use of the PANNORAMIC MIDI II digital slide scanner and thank P. Vági and L. Barna for their assistance in image acquisition.

## AUTHOR CONTRIBUTIONS

Conceptualization: VT, PP, ÁO, TFF, GN; Neurolucida reconstructions: VT, PP, TZ, KZ; Electron microscopy: VT, ZB, TZ; Data analyses: VT, PP, ÁO, ZB, KZ; Tissue sampling: ZM, PG; Antibody generation: MW; Funding acquisition: TFF, GN; Supervision: ZM, GN; Writing—original draft: VT, PP, ÁO, GN; Writing—review & editing: VT, PP, ÁO, ZB, TZ, KZ, MW, ZM, PG, TFF, GN.

## Extended data

**Extended Data Table 1:** Densities and absolute numbers of interneurons across layers in consecutive blocks of a single hippocampal hemisphere in subjects SKO24, 25, 27, and 31.

**Subject: SKO24, Immunohistochemistry: PV**
| Block | 1 from 0 um - (Head) |  |  | 2 from 8500 um - |  |  | 3 from 20540 um - |  |  | 4 from 25480 um - |  |  | Total |  |  |
| --- | --- | --- | --- | --- | --- | --- | --- | --- | --- | --- | --- | --- | --- | --- | --- |
| SKO24 PV | Volume [mm<sup>3</sup>] | Density [cell/mm<sup>3</sup>] | Abs. No | Volume [mm<sup>3</sup>] | Density [cell/mm<sup>3</sup>] | Abs. No | Volume [mm<sup>3</sup>] | Density [cell/mm<sup>3</sup>] | Abs. No | Volume [mm<sup>3</sup>] | Density [cell/mm<sup>3</sup>] | Abs. No | Volume [mm<sup>3</sup>] | Density [cell/mm<sup>3</sup>] | Abs. No |
| CA1 ori | 23.8 | 11.7 | 278.0 | 11.8 | 186.5 | 2208.7 | 7.2 | 287.6 | 2066.1 | 1.4 | 375.6 | 542.1 | 44.3 | 115.1 | 5094.9 |
| CA1pyr | 54.7 | 170.8 | 9338.6 | 31.1 | 176.2 | 5480.8 | 22.7 | 339.2 | 7695.9 | 4.2 | 430.1 | 1804.3 | 112.7 | 215.9 | 24319.5 |
| CA1 rad | 28.1 | 8.9 | 251.4 | 15.3 | 18.8 | 286.5 | 10.8 | 62.4 | 676.1 | 2.4 | 25.5 | 62.1 | 56.7 | 22.5 | 1276.1 |
| CA1 lm | 32.8 | 0.0 | 0.0 | 18.5 | 5.3 | 98.3 | 15.9 | 27.7 | 441.7 | 1.0 | 0.0 | 0.0 | 68.3 | 7.9 | 540.0 |
| CA2 ori | 5.7 | 0.0 | 0.0 | 5.9 | 234.4 | 1379.5 | 2.6 | 60.0 | 158.1 | 0.4 | 162.3 | 72.0 | 14.7 | 109.8 | 1609.6 |
| CA2 pyr | 9.2 | 71.4 | 656.8 | 9.5 | 117.0 | 1109.6 | 4.7 | 192.8 | 914.9 | 0.9 | 251.6 | 216.0 | 24.3 | 119.3 | 2897.2 |
| CA2 rad | 5.1 | 0.0 | 0.0 | 6.1 | 0.0 | 0.0 | 2.0 | 0.0 | 0.0 | 0.5 | 0.0 | 0.0 | 13.8 | 0.0 | 0.0 |
| CA2 lm | 2.6 | 0.0 | 0.0 | 3.0 | 0.0 | 0.0 | 2.8 | 0.0 | 0.0 | 0.9 | 0.0 | 0.0 | 9.3 | 0.0 | 0.0 |
| CA3 ori | 4.6 | 9.8 | 45.6 | 10.8 | 16.2 | 175.3 | 5.3 | 61.5 | 328.8 | 1.5 | 23.4 | 36.0 | 22.3 | 26.3 | 585.7 |
| CA3 pyr | 19.7 | 37.1 | 730.6 | 40.1 | 59.6 | 2390.9 | 20.7 | 100.7 | 2083.6 | 2.9 | 125.0 | 360.0 | 83.4 | 66.7 | 5565.1 |
| CA3 rad | 1.2 | 0.0 | 0.0 | 5.0 | 0.0 | 0.0 | 2.8 | 37.5 | 106.2 | 0.6 | 0.0 | 0.0 | 9.6 | 11.0 | 106.2 |
| CA3 lm | 0.6 | 0.0 | 0.0 | 1.3 | 0.0 | 0.0 | 0.9 | 0.0 | 0.0 | 0.2 | 0.0 | 0.0 | 3.1 | 0.0 | 0.0 |
| DG mol | 7.5 | 0.0 | 0.0 | 26.6 | 4.2 | 112.0 | 17.3 | 14.1 | 245.0 | 1.3 | 0.0 | 0.0 | 52.8 | 6.8 | 357.0 |
| DG gra | 4.2 | 116.7 | 494.4 | 15.0 | 83.1 | 1247.7 | 6.5 | 293.1 | 1915.8 | 0.5 | 66.8 | 36.0 | 26.3 | 140.4 | 3693.8 |
| DG hil | 4.5 | 11.4 | 51.8 | 11.9 | 87.7 | 1042.0 | 4.4 | 289.2 | 1259.8 | 0.8 | 328.0 | 252.0 | 21.6 | 120.9 | 2605.6 |
| Sum of all |  |  |  |  |  |  |  |  |  |  |  |  | 562.9 |  | 48650.9 |

**Subject: SKO24, Immunohistochemistry: SOM**
| Block | 1 from 0 um - (Head) |  |  | 2 from 8500 um - |  |  | 3 from 20540 um - |  |  | 4 from 25480 um - |  |  | Total |  |  |
| --- | --- | --- | --- | --- | --- | --- | --- | --- | --- | --- | --- | --- | --- | --- | --- |
| SKO24 SOM | Volume [mm<sup>3</sup>] | Density [cell/mm<sup>3</sup>] | Abs. No | Volume [mm<sup>3</sup>] | Density [cell/mm<sup>3</sup>] | Abs. No | Volume [mm<sup>3</sup>] | Density [cell/mm<sup>3</sup>] | Abs. No | Volume [mm<sup>3</sup>] | Density [cell/mm<sup>3</sup>] | Abs. No | Volume [mm<sup>3</sup>] | Density [cell/mm<sup>3</sup>] | Abs. No |
| CA1 ori | 23.8 | 410.8 | 9775.2 | 11.8 | 514.5 | 6092.9 | 7.2 | 587.5 | 4220.2 | 1.4 | 101.8 | 146.9 | 44.3 | 457.1 | 20235.2 |
| CA1pyr | 54.7 | 336.8 | 18413.7 | 31.1 | 362.5 | 11273.1 | 22.7 | 404.9 | 9186.5 | 4.2 | 367.8 | 1542.8 | 112.7 | 358.8 | 40416.1 |
| CA1 rad | 28.1 | 90.0 | 2530.4 | 15.3 | 48.3 | 737.5 | 10.8 | 91.2 | 988.3 | 2.4 | 50.2 | 122.5 | 56.7 | 77.3 | 4378.7 |
| CA1 lm | 32.8 | 15.6 | 510.3 | 18.5 | 8.9 | 164.6 | 15.9 | 33.1 | 527.4 | 1.0 | 0.0 | 0.0 | 68.3 | 17.6 | 1202.3 |
| CA2 ori | 5.7 | 577.1 | 3289.5 | 5.9 | 362.6 | 2133.8 | 2.6 | 173.2 | 456.1 | 0.4 | 506.4 | 224.7 | 14.7 | 416.3 | 6104.1 |
| CA2 pyr | 9.2 | 336.9 | 3099.3 | 9.5 | 109.6 | 1039.5 | 4.7 | 146.9 | 696.9 | 0.9 | 70.5 | 60.5 | 24.3 | 201.6 | 4896.3 |
| CA2 rad | 5.1 | 212.0 | 1080.7 | 6.1 | 82.0 | 501.5 | 2.0 | 32.0 | 64.9 | 0.5 | 0.0 | 0.0 | 13.8 | 119.8 | 1647.1 |
| CA2 lm | 2.6 | 0.0 | 0.0 | 3.0 | 21.1 | 64.1 | 2.8 | 0.0 | 0.0 | 0.9 | 0.0 | 0.0 | 9.3 | 6.9 | 64.1 |
| CA3 ori | 4.6 | 322.1 | 1493.0 | 10.8 | 358.0 | 3862.1 | 5.3 | 370.7 | 1980.6 | 1.5 | 418.0 | 642.1 | 22.3 | 357.7 | 7977.8 |
| CA3 pyr | 19.7 | 493.2 | 9704.3 | 40.1 | 502.6 | 20177.6 | 20.7 | 557.7 | 11535.0 | 2.9 | 727.7 | 2095.7 | 83.4 | 521.8 | 43512.6 |
| CA3 rad | 1.2 | 86.9 | 103.4 | 5.0 | 74.3 | 370.3 | 2.8 | 249.5 | 707.0 | 0.6 | 0.0 | 0.0 | 9.6 | 122.7 | 1180.8 |
| CA3 lm | 0.6 | 0.0 | 0.0 | 1.3 | 0.0 | 0.0 | 0.9 | 0.0 | 0.0 | 0.2 | 0.0 | 0.0 | 3.1 | 0.0 | 0.0 |
| DG mol | 7.5 | 0.0 | 0.0 | 26.6 | 12.8 | 342.2 | 17.3 | 26.7 | 461.8 | 1.3 | 14.2 | 19.0 | 52.8 | 15.6 | 823.0 |
| DG gra | 4.2 | 34.3 | 145.5 | 15.0 | 92.5 | 1388.7 | 6.5 | 73.5 | 480.5 | 0.5 | 0.0 | 0.0 | 26.3 | 76.6 | 2014.7 |
| DG hil | 4.5 | 543.9 | 2473.3 | 11.9 | 766.1 | 9105.1 | 4.4 | 1266.8 | 5518.3 | 0.8 | 1359.5 | 1044.6 | 21.6 | 841.6 | 18141.3 |
| Sum of all |  |  |  |  |  |  |  |  |  |  |  |  | 562.9 |  | 152594 |

**Subject: SKO24, Immunohistochemistry: CR**
| Block | 1 from 0 um - (Head) |  |  | 2 from 8500 um - |  |  | 3 from 20540 um - |  |  | 4 from 25480 um - |  |  | Total |  |  |
| --- | --- | --- | --- | --- | --- | --- | --- | --- | --- | --- | --- | --- | --- | --- | --- |
| SKO24 CR | Volume [mm<sup>3</sup>] | Density [cell/mm<sup>3</sup>] | Abs. No | Volume [mm<sup>3</sup>] | Density [cell/mm<sup>3</sup>] | Abs. No | Volume [mm<sup>3</sup>] | Density [cell/mm<sup>3</sup>] | Abs. No | Volume [mm<sup>3</sup>] | Density [cell/mm<sup>3</sup>] | Abs. No | Volume [mm<sup>3</sup>] | Density [cell/mm<sup>3</sup>] | Abs. No |
| CA1 ori | 23.8 | 128.6 | 3060.5 | 11.8 | 246.2 | 2915.5 | 7.2 | 151.9 | 1091.2 | 1.4 | 72.8 | 105.1 | 44.3 | 162.0 | 7172.2 |
| CA1pyr | 54.7 | 200.1 | 10935.9 | 31.1 | 261.0 | 8119.2 | 22.7 | 312.5 | 7090.3 | 4.2 | 423.3 | 1775.6 | 112.7 | 247.9 | 27920.9 |
| CA1 rad | 28.1 | 305.1 | 8581.1 | 15.3 | 528.6 | 8071.0 | 10.8 | 755.8 | 8186.0 | 2.4 | 1841.3 | 4490.8 | 56.7 | 517.6 | 29329.0 |
| CA1 lm | 32.8 | 467.6 | 15330.3 | 18.5 | 438.1 | 8102.8 | 15.9 | 697.2 | 11119.6 | 1.0 | 1953.7 | 2014.3 | 68.3 | 535.7 | 36567.0 |
| CA2 ori | 5.7 | 287.3 | 1637.6 | 5.9 | 122.2 | 719.2 | 2.6 | 38.7 | 101.9 | 0.4 | 237.5 | 105.4 | 14.7 | 174.9 | 2564.1 |
| CA2 pyr | 9.2 | 237.9 | 2188.1 | 9.5 | 144.9 | 1374.4 | 4.7 | 25.3 | 120.0 | 0.9 | 37.1 | 31.8 | 24.3 | 152.9 | 3714.3 |
| CA2 rad | 5.1 | 271.4 | 1383.9 | 6.1 | 40.6 | 248.4 | 2.0 | 141.5 | 286.8 | 0.5 | 233.7 | 119.7 | 13.8 | 148.3 | 2038.8 |
| CA2 lm | 2.6 | 0.0 | 0.0 | 3.0 | 232.6 | 707.4 | 2.8 | 182.7 | 508.2 | 0.9 | 292.1 | 259.0 | 9.3 | 157.8 | 1474.6 |
| CA3 ori | 4.6 | 142.9 | 662.1 | 10.8 | 203.8 | 2198.4 | 5.3 | 274.0 | 1463.8 | 1.5 | 332.2 | 510.3 | 22.3 | 216.8 | 4834.5 |
| CA3 pyr | 19.7 | 221.0 | 4348.0 | 40.1 | 155.1 | 6225.9 | 20.7 | 197.9 | 4093.0 | 2.9 | 351.5 | 1012.5 | 83.4 | 188.0 | 15679.4 |
| CA3 rad | 1.2 | 335.8 | 399.7 | 5.0 | 176.3 | 878.1 | 2.8 | 503.7 | 1427.5 | 0.6 | 198.2 | 121.6 | 9.6 | 293.9 | 2826.9 |
| CA3 lm | 0.6 | 0.0 | 0.0 | 1.3 | 113.5 | 153.1 | 0.9 | 95.2 | 88.7 | 0.2 | 233.7 | 44.5 | 3.1 | 93.0 | 286.3 |
| DG mol | 7.5 | 396.8 | 2956.3 | 26.6 | 428.4 | 11415.0 | 17.3 | 372.4 | 6451.8 | 1.3 | 521.7 | 697.4 | 52.8 | 407.9 | 21520.6 |
| DG gra | 4.2 | 1042.8 | 4418.7 | 15.0 | 1184.7 | 17777.1 | 6.5 | 1332.0 | 8707.9 | 0.5 | 1733.0 | 933.7 | 26.3 | 1209.7 | 31837.5 |
| DG hil | 4.5 | 98.6 | 448.2 | 11.9 | 913.8 | 10860.0 | 4.4 | 1234.7 | 5378.7 | 0.8 | 1629.4 | 1252.0 | 21.6 | 832.2 | 17938.9 |
| Sum of all |  |  |  |  |  |  |  |  |  |  |  |  | 562.9 |  | 205705.1 |

Subject: SKO25, Immunohistochemistry: PV
| Block | 1 from 0 um - (Head) |  |  | 2 from 7000 um - |  |  | 3 from 18240 um - |  |  | 4 from 29500 um - |  |  | 5 from 38900 um - |  |  | Total |  |  |
| --- | --- | --- | --- | --- | --- | --- | --- | --- | --- | --- | --- | --- | --- | --- | --- | --- | --- | --- |
| SKO25 PV | Volume [mm<sup>3</sup>] | Density [cell/mm<sup>3</sup>] | Abs. No | Volume [mm<sup>3</sup>] | Density [cell/mm<sup>3</sup>] | Abs. No | Volume [mm<sup>3</sup>] | Density [cell/mm<sup>3</sup>] | Abs. No | Volume [mm<sup>3</sup>] | Density [cell/mm<sup>3</sup>] | Abs. No | Volume [mm<sup>3</sup>] | Density [cell/mm<sup>3</sup>] | Abs. No | Volume [mm<sup>3</sup>] | Density [cell/mm<sup>3</sup>] | Abs. No |
| CA1 ori | 19.2 | 118.1 | 2263.7 | 8.0 | 246.2 | 1958.6 | 8.3 | 274.0 | 2284.5 | 2.0 | 169.3 | 338.5 | 1.7 | 115.0 | 195.5 | 39.2 | 179.8 | 7040.7 |
| CA1pyr | 42.4 | 37.0 | 1570.3 | 21.8 | 129.0 | 2806.6 | 15.0 | 181.9 | 2722.2 | 5.1 | 157.2 | 804.4 | 2.5 | 180.6 | 455.7 | 86.8 | 96.3 | 8359.3 |
| CA1 rad | 24.7 | 2.0 | 48.9 | 17.1 | 5.4 | 92.1 | 10.6 | 12.2 | 129.7 | 5.7 | 25.0 | 142.2 | 2.0 | 0.0 | 0.0 | 60.0 | 6.9 | 413.0 |
| CA1 lm | 11.2 | 11.1 | 125.0 | 14.1 | 13.9 | 195.7 | 8.3 | 47.0 | 388.2 | 4.4 | 0.0 | 0.0 | 1.4 | 43.3 | 59.8 | 39.3 | 19.6 | 768.6 |
| CA2 ori | 3.2 | 38.8 | 125.0 | 4.8 | 78.5 | 378.5 | 3.2 | 139.6 | 443.0 | 2.2 | 0.0 | 0.0 | 2.2 | 88.7 | 191.8 | 15.6 | 72.9 | 1138.3 |
| CA2 pyr | 4.3 | 58.1 | 250.0 | 6.2 | 123.3 | 760.2 | 4.4 | 165.9 | 724.6 | 2.7 | 141.1 | 375.1 | 2.4 | 195.0 | 459.0 | 19.9 | 129.4 | 2568.8 |
| CA2 rad | 7.2 | 0.0 | 0.0 | 9.7 | 0.0 | 0.0 | 5.7 | 0.0 | 0.0 | 2.9 | 0.0 | 0.0 | 2.7 | 0.0 | 0.0 | 28.1 | 0.0 | 0.0 |
| CA2 lm | 4.5 | 18.3 | 83.3 | 4.4 | 18.1 | 80.2 | 3.6 | 0.0 | 0.0 | 1.6 | 0.0 | 0.0 | 0.5 | 0.0 | 0.0 | 14.6 | 11.2 | 163.5 |
| CA3 ori | 0.0 | 0.0 | 0.0 | 13.4 | 8.1 | 107.7 | 16.6 | 17.6 | 293.6 | 4.9 | 55.5 | 270.5 | 5.2 | 10.3 | 53.6 | 40.1 | 18.1 | 725.4 |
| CA3 pyr | 29.6 | 22.5 | 666.7 | 48.5 | 28.6 | 1384.3 | 37.2 | 58.8 | 2187.1 | 25.4 | 97.1 | 2464.4 | 17.6 | 136.2 | 2394.4 | 158.2 | 57.5 | 9096.9 |
| CA3 rad | 0.0 | 0.0 | 0.0 | 9.3 | 28.9 | 268.8 | 6.4 | 0.0 | 0.0 | 2.0 | 0.0 | 0.0 | 0.8 | 0.0 | 0.0 | 18.5 | 14.5 | 268.8 |
| CA3 lm | 0.0 | 0.0 | 0.0 | 2.2 | 45.1 | 101.3 | 1.8 | 0.0 | 0.0 | 0.6 | 0.0 | 0.0 | 0.1 | 0.0 | 0.0 | 4.7 | 21.4 | 101.3 |
| DG mol | 16.0 | 10.4 | 166.7 | 27.9 | 5.4 | 151.9 | 30.5 | 16.1 | 490.9 | 18.5 | 31.7 | 587.5 | 12.9 | 36.7 | 473.5 | 105.8 | 17.7 | 1870.5 |
| DG gra | 6.5 | 57.3 | 375.0 | 9.5 | 37.2 | 352.6 | 13.5 | 81.8 | 1102.4 | 6.6 | 82.6 | 545.3 | 4.6 | 149.5 | 688.3 | 40.7 | 75.2 | 3063.5 |
| DG hil | 6.0 | 76.8 | 458.3 | 11.3 | 63.9 | 719.5 | 15.3 | 71.8 | 1101.8 | 5.9 | 169.5 | 1008.0 | 7.3 | 188.9 | 1373.0 | 45.8 | 101.8 | 4660.6 |
| Sum of all |  |  |  |  |  |  |  |  |  |  |  |  |  |  |  | 717.4 |  | 40239.2 |

Subject: SKO25, Immunohistochemistry: SOM
| Block | 1 from 0 um - (Head) |  |  | 2 from 7000 um - |  |  | 3 from 18240 um - |  |  | 4 from 29500 um - |  |  | 5 from 38900 um - |  |  | Total |  |  |
| --- | --- | --- | --- | --- | --- | --- | --- | --- | --- | --- | --- | --- | --- | --- | --- | --- | --- | --- |
| SKO25 SOM | Volume [mm<sup>3</sup>] | Density [cell/mm<sup>3</sup>] | Abs. No | Volume [mm<sup>3</sup>] | Density [cell/mm<sup>3</sup>] | Abs. No | Volume [mm<sup>3</sup>] | Density [cell/mm<sup>3</sup>] | Abs. No | Volume [mm<sup>3</sup>] | Density [cell/mm<sup>3</sup>] | Abs. No | Volume [mm<sup>3</sup>] | Density [cell/mm<sup>3</sup>] | Abs. No | Volume [mm<sup>3</sup>] | Density [cell/mm<sup>3</sup>] | Abs. No |
| CA1 ori | 19.2 | 243.5 | 4668.0 | 8.0 | 127.8 | 1016.7 | 8.3 | 375.6 | 3131.2 | 2.0 | 402.3 | 804.1 | 1.7 | 42.1 | 71.6 | 39.2 | 247.5 | 9691.5 |
| CA1pyr | 42.4 | 65.9 | 2794.8 | 21.8 | 260.2 | 5661.2 | 15.0 | 304.9 | 4564.0 | 5.1 | 257.0 | 1315.2 | 2.5 | 85.6 | 215.9 | 86.8 | 167.7 | 14551.1 |
| CA1 rad | 24.7 | 20.6 | 507.9 | 17.1 | 28.7 | 490.1 | 10.6 | 59.0 | 625.6 | 5.7 | 21.9 | 124.5 | 2.0 | 0.0 | 0.0 | 60.0 | 29.1 | 1748.1 |
| CA1 lm | 11.2 | 0.0 | 0.0 | 14.1 | 6.0 | 85.1 | 8.3 | 0.0 | 0.0 | 4.4 | 0.0 | 0.0 | 1.4 | 0.0 | 0.0 | 39.3 | 2.2 | 85.1 |
| CA2 ori | 3.2 | 110.1 | 354.4 | 4.8 | 121.2 | 584.8 | 3.2 | 137.7 | 436.9 | 2.2 | 101.2 | 227.1 | 2.2 | 468.9 | 1013.7 | 15.6 | 167.5 | 2617.0 |
| CA2 pyr | 4.3 | 111.2 | 478.8 | 6.2 | 182.7 | 1126.5 | 4.4 | 185.5 | 809.9 | 2.7 | 226.9 | 603.4 | 2.4 | 94.8 | 223.0 | 19.9 | 163.3 | 3241.7 |
| CA2 rad | 7.2 | 36.9 | 264.5 | 9.7 | 9.3 | 90.1 | 5.7 | 0.0 | 0.0 | 2.9 | 0.0 | 0.0 | 2.7 | 50.3 | 134.1 | 28.1 | 17.4 | 488.7 |
| CA2 lm | 4.5 | 5.0 | 22.7 | 4.4 | 0.0 | 0.0 | 3.6 | 0.0 | 0.0 | 1.6 | 0.0 | 0.0 | 0.5 | 0.0 | 0.0 | 14.6 | 1.5 | 22.7 |
| CA3 ori | 0.0 | 0.0 | 0.0 | 13.4 | 322.6 | 4314.1 | 16.6 | 282.4 | 4700.5 | 4.9 | 231.2 | 1127.7 | 5.2 | 371.2 | 1932.2 | 40.1 | 301.1 | 12074.4 |
| CA3 pyr | 29.6 | 480.4 | 14203.5 | 48.5 | 377.9 | 18321.3 | 37.2 | 296.2 | 11012.0 | 25.4 | 263.2 | 6682.9 | 17.6 | 375.6 | 6604.5 | 158.2 | 359.2 | 56824.2 |
| CA3 rad | 0.0 | 0.0 | 0.0 | 9.3 | 120.5 | 1119.5 | 6.4 | 64.9 | 416.0 | 2.0 | 19.6 | 38.9 | 0.8 | 517.4 | 424.3 | 18.5 | 108.0 | 1998.7 |
| CA3 lm | 0.0 | 0.0 | 0.0 | 2.2 | 0.0 | 0.0 | 1.8 | 0.0 | 0.0 | 0.6 | 0.0 | 0.0 | 0.1 | 0.0 | 0.0 | 4.7 | 0.0 | 0.0 |
| DG mol | 16.0 | 5.2 | 83.1 | 27.9 | 2.6 | 73.8 | 30.5 | 1.9 | 59.1 | 18.5 | 0.0 | 0.0 | 12.9 | 0.0 | 0.0 | 105.8 | 2.0 | 216.1 |
| DG gra | 6.5 | 26.2 | 171.6 | 9.5 | 31.2 | 296.2 | 13.5 | 28.7 | 386.5 | 6.6 | 33.7 | 222.4 | 4.6 | 11.3 | 52.1 | 40.7 | 27.7 | 1128.8 |
| DG hil | 6.0 | 276.8 | 1650.8 | 11.3 | 439.4 | 4944.0 | 15.3 | 477.3 | 7323.5 | 5.9 | 597.2 | 3551.8 | 7.3 | 1142.6 | 8303.1 | 45.8 | 563.1 | 25773.2 |
| Sum of all |  |  |  |  |  |  |  |  |  |  |  |  |  |  |  | 717.4 |  | 130461.2 |

Subject: SKO25, Immunohistochemistry: CR
| Block | 1 from 0 um - (Head) |  |  | 2 from 7000 um - |  |  | 3 from 18240 um - |  |  | 4 from 29500 um - |  |  | 5 from 38900 um - |  |  | Total |  |  |
| --- | --- | --- | --- | --- | --- | --- | --- | --- | --- | --- | --- | --- | --- | --- | --- | --- | --- | --- |
| SKO25 CR | Volume [mm<sup>3</sup>] | Density [cell/mm<sup>3</sup>] | Abs. No | Volume [mm<sup>3</sup>] | Density [cell/mm<sup>3</sup>] | Abs. No | Volume [mm<sup>3</sup>] | Density [cell/mm<sup>3</sup>] | Abs. No | Volume [mm<sup>3</sup>] | Density [cell/mm<sup>3</sup>] | Abs. No | Volume [mm<sup>3</sup>] | Density [cell/mm<sup>3</sup>] | Abs. No | Volume [mm<sup>3</sup>] | Density [cell/mm<sup>3</sup>] | Abs. No |
| CA1 ori | 19.2 | 40.7 | 780.3 | 8.0 | 70.6 | 561.7 | 8.3 | 101.9 | 849.9 | 2.0 | 130.9 | 261.7 | 1.7 | 159.2 | 270.6 | 39.2 | 69.6 | 2724.1 |
| CA1pyr | 42.4 | 226.4 | 9606.3 | 21.8 | 285.9 | 6219.5 | 15.0 | 191.9 | 2872.5 | 5.1 | 217.4 | 1112.8 | 2.5 | 69.1 | 174.5 | 86.8 | 230.3 | 19985.5 |
| CA1 rad | 24.7 | 688.5 | 16987.8 | 17.1 | 833.9 | 14223.9 | 10.6 | 1138.6 | 12062.0 | 5.7 | 1024.5 | 5823.8 | 2.0 | 294.6 | 596.5 | 60.0 | 827.8 | 49694.0 |
| CA1 lm | 11.2 | 484.2 | 5434.8 | 14.1 | 547.9 | 7717.7 | 8.3 | 420.4 | 3474.4 | 4.4 | 309.4 | 1348.4 | 1.4 | 123.5 | 170.5 | 39.3 | 461.6 | 18145.8 |
| CA2 ori | 3.2 | 37.2 | 119.8 | 4.8 | 53.9 | 259.9 | 3.2 | 55.6 | 176.5 | 2.2 | 120.4 | 270.1 | 2.2 | 181.8 | 392.9 | 15.6 | 78.0 | 1219.2 |
| CA2 pyr | 4.3 | 150.4 | 647.7 | 6.2 | 96.6 | 595.5 | 4.4 | 90.9 | 396.8 | 2.7 | 90.9 | 241.7 | 2.4 | 109.1 | 256.8 | 19.9 | 107.7 | 2138.4 |
| CA2 rad | 7.2 | 228.7 | 1639.0 | 9.7 | 61.4 | 596.3 | 5.7 | 153.2 | 869.6 | 2.9 | 192.4 | 558.9 | 2.7 | 276.2 | 736.8 | 28.1 | 156.5 | 4400.5 |
| CA2 lm | 4.5 | 509.5 | 2315.4 | 4.4 | 177.4 | 785.1 | 3.6 | 215.9 | 774.5 | 1.6 | 190.2 | 296.2 | 0.5 | 71.1 | 36.4 | 14.6 | 287.6 | 4207.6 |
| CA3 ori | 0.0 | 0.0 | 0.0 | 13.4 | 75.0 | 1003.0 | 16.6 | 148.7 | 2474.7 | 4.9 | 99.8 | 486.9 | 5.2 | 335.1 | 1744.3 | 40.1 | 142.4 | 5708.9 |
| CA3 pyr | 29.6 | 96.0 | 2837.5 | 48.5 | 75.4 | 3656.9 | 37.2 | 166.3 | 6183.0 | 25.4 | 170.4 | 4326.1 | 17.6 | 261.3 | 4595.3 | 158.2 | 136.5 | 21598.8 |
| CA3 rad | 0.0 | 0.0 | 0.0 | 9.3 | 174.9 | 1625.0 | 6.4 | 288.1 | 1845.3 | 2.0 | 153.3 | 304.8 | 0.8 | 818.2 | 671.0 | 18.5 | 240.3 | 4446.1 |
| CA3 lm | 0.0 | 0.0 | 0.0 | 2.2 | 84.5 | 189.8 | 1.8 | 484.7 | 879.8 | 0.6 | 372.9 | 227.9 | 0.1 | 935.1 | 56.9 | 4.7 | 286.1 | 1354.3 |
| DG mol | 16.0 | 173.2 | 2769.1 | 27.9 | 292.9 | 8185.1 | 30.5 | 297.7 | 9072.4 | 18.5 | 308.3 | 5710.5 | 12.9 | 331.5 | 4272.5 | 105.8 | 283.6 | 30009.6 |
| DG gra | 6.5 | 315.8 | 2065.1 | 9.5 | 345.7 | 3278.4 | 13.5 | 993.7 | 13396.7 | 6.6 | 767.0 | 5063.3 | 4.6 | 925.3 | 4261.2 | 40.7 | 689.3 | 28064.7 |
| DG hil | 6.0 | 303.6 | 1811.1 | 11.3 | 531.1 | 5976.2 | 15.3 | 1150.0 | 17644.5 | 5.9 | 1086.5 | 6462.0 | 7.3 | 943.6 | 6857.0 | 45.8 | 846.6 | 38750.7 |
| Sum of all |  |  |  |  |  |  |  |  |  |  |  |  |  |  |  | 717.4 |  | 232448.0 |

Subject: SKO27, Immunohistochemistry: PV
| Block | 1 from 0 um - (Head) |  |  | 2 from 3980 um - |  |  | 3 from 12580 um - |  |  | 4 from 22180 um - |  |  | 5 from 31960 um - |  |  | Total |  |  |
| --- | --- | --- | --- | --- | --- | --- | --- | --- | --- | --- | --- | --- | --- | --- | --- | --- | --- | --- |
| SKO27 PV | Volume [mm<sup>3</sup>] | Density [cell/mm<sup>3</sup>] | Abs. No | Volume [mm<sup>3</sup>] | Density [cell/mm<sup>3</sup>] | Abs. No | Volume [mm<sup>3</sup>] | Density [cell/mm<sup>3</sup>] | Abs. No | Volume [mm<sup>3</sup>] | Density [cell/mm<sup>3</sup>] | Abs. No | Volume [mm<sup>3</sup>] | Density [cell/mm<sup>3</sup>] | Abs. No | Volume [mm<sup>3</sup>] | Density [cell/mm<sup>3</sup>] | Abs. No |
| CA1 ori | 28.7 | 66.8 | 1918.9 | 13.2 | 105.8 | 1392.7 | 10.4 | 135.2 | 1408.9 | 13.0 | 177.5 | 2308.7 | 9.8 | 322.9 | 3151.1 | 75.1 | 135.6 | 10180.4 |
| CA1pyr | 68.9 | 54.9 | 3786.5 | 42.3 | 74.1 | 3136.2 | 19.0 | 111.1 | 2110.4 | 20.1 | 110.4 | 2214.8 | 27.5 | 160.8 | 4423.3 | 177.8 | 88.1 | 15671.2 |
| CA1 rad | 31.1 | 0.0 | 0.0 | 17.1 | 4.8 | 81.6 | 19.6 | 59.1 | 1161.6 | 15.4 | 24.6 | 378.3 | 14.8 | 51.1 | 756.1 | 98.0 | 24.3 | 2377.6 |
| CA1 Im | 53.3 | 3.1 | 163.7 | 16.9 | 2.7 | 45.6 | 7.3 | 57.4 | 418.8 | 10.2 | 39.5 | 403.7 | 11.2 | 334.6 | 3731.3 | 98.9 | 48.2 | 4763.0 |
| CA2 ori | 5.9 | 41.5 | 244.7 | 4.2 | 95.4 | 401.0 | 2.0 | 149.1 | 300.8 | 3.5 | 137.7 | 488.5 | 0.9 | 100.7 | 89.2 | 16.6 | 92.1 | 1524.3 |
| CA2 pyr | 4.3 | 94.3 | 402.6 | 5.3 | 88.1 | 463.9 | 2.3 | 235.3 | 541.1 | 2.9 | 171.1 | 502.0 | 1.6 | 216.3 | 356.7 | 16.4 | 138.0 | 2266.2 |
| CA2 rad | 5.2 | 0.0 | 0.0 | 7.7 | 0.0 | 0.0 | 2.9 | 0.0 | 0.0 | 2.5 | 0.0 | 0.0 | 1.5 | 0.0 | 0.0 | 19.9 | 0.0 | 0.0 |
| CA2 Im | 3.4 | 8.1 | 27.8 | 3.9 | 0.0 | 0.0 | 1.5 | 0.0 | 0.0 | 2.3 | 0.0 | 0.0 | 2.0 | 21.8 | 42.7 | 13.2 | 5.4 | 70.6 |
| CA3 ori | 7.1 | 6.9 | 49.6 | 14.9 | 24.5 | 365.4 | 14.1 | 23.7 | 335.1 | 8.9 | 108.0 | 959.8 | 10.7 | 60.8 | 649.5 | 55.8 | 42.3 | 2359.4 |
| CA3 pyr | 27.3 | 49.1 | 1340.2 | 77.7 | 31.7 | 2461.6 | 32.5 | 40.8 | 1326.4 | 28.7 | 55.3 | 1584.8 | 25.0 | 117.0 | 2927.9 | 191.2 | 50.4 | 9640.9 |
| CA3 rad | 10.3 | 5.2 | 53.4 | 9.9 | 3.5 | 34.2 | 8.0 | 0.0 | 0.0 | 3.7 | 0.0 | 0.0 | 3.4 | 13.2 | 44.4 | 35.2 | 3.8 | 132.0 |
| CA3 Im | 2.4 | 10.5 | 24.8 | 4.4 | 8.2 | 35.8 | 1.9 | 0.0 | 0.0 | 1.2 | 0.0 | 0.0 | 1.7 | 24.9 | 42.2 | 11.6 | 8.9 | 102.8 |
| DG mol | 30.4 | 12.3 | 375.2 | 41.0 | 2.5 | 104.1 | 32.8 | 15.1 | 496.9 | 26.0 | 8.5 | 220.4 | 27.9 | 9.4 | 263.1 | 158.2 | 9.2 | 1459.7 |
| DG gra | 9.5 | 67.3 | 638.8 | 44.6 | 6.3 | 279.6 | 10.1 | 20.9 | 210.4 | 11.4 | 165.8 | 1893.2 | 7.5 | 196.8 | 1479.8 | 83.1 | 54.2 | 4501.8 |
| DG hil | 10.7 | 18.2 | 194.4 | 15.1 | 33.3 | 503.3 | 9.6 | 67.8 | 651.9 | 12.1 | 102.2 | 1238.7 | 8.1 | 135.5 | 1095.1 | 55.7 | 66.2 | 3683.4 |
| Sum of all |  |  |  |  |  |  |  |  |  |  |  |  |  |  |  | 1106.5 |  | 58733.1 |

Subject: SKO27, Immunohistochemistry: SOM
| Block | 1 from 0 um - (Head) |  |  | 2 from 3980 um - |  |  | 3 from 12580 um - |  |  | 4 from 22180 um - |  |  | 5 from 31960 um - |  |  | Total |  |  |
| --- | --- | --- | --- | --- | --- | --- | --- | --- | --- | --- | --- | --- | --- | --- | --- | --- | --- | --- |
| SKO27 SOM | Volume [mm<sup>3</sup>] | Density [cell/mm<sup>3</sup>] | Abs. No | Volume [mm<sup>3</sup>] | Density [cell/mm<sup>3</sup>] | Abs. No | Volume [mm<sup>3</sup>] | Density [cell/mm<sup>3</sup>] | Abs. No | Volume [mm<sup>3</sup>] | Density [cell/mm<sup>3</sup>] | Abs. No | Volume [mm<sup>3</sup>] | Density [cell/mm<sup>3</sup>] | Abs. No | Volume [mm<sup>3</sup>] | Density [cell/mm<sup>3</sup>] | Abs. No |
| CA1 ori | 28.7 | 281.8 | 8094.2 | 13.2 | 35.2 | 463.9 | 10.4 | 157.2 | 1638.3 | 13.0 | 135.9 | 1767.9 | 9.8 | 25.2 | 246.2 | 75.1 | 162.7 | 12210.6 |
| CA1pyr | 68.9 | 55.2 | 3801.7 | 42.3 | 5.7 | 241.7 | 19.0 | 20.1 | 382.7 | 20.1 | 22.0 | 441.5 | 27.5 | 31.2 | 857.2 | 177.8 | 32.2 | 5724.8 |
| CA1 rad | 31.1 | 37.6 | 1169.5 | 17.1 | 0.0 | 0.0 | 19.6 | 7.0 | 138.3 | 15.4 | 14.0 | 215.5 | 14.8 | 0.0 | 0.0 | 98.0 | 15.5 | 1523.4 |
| CA1 Im | 53.3 | 3.9 | 209.0 | 16.9 | 0.0 | 0.0 | 7.3 | 0.0 | 0.0 | 10.2 | 0.0 | 0.0 | 11.2 | 0.0 | 0.0 | 98.9 | 2.1 | 209.0 |
| CA2 ori | 5.9 | 394.4 | 2327.3 | 4.2 | 40.9 | 171.9 | 2.0 | 90.7 | 183.0 | 3.5 | 156.0 | 553.7 | 0.9 | 115.4 | 102.2 | 16.6 | 201.6 | 3338.1 |
| CA2 pyr | 4.3 | 61.7 | 263.4 | 5.3 | 21.8 | 114.8 | 2.3 | 65.7 | 151.1 | 2.9 | 40.4 | 118.6 | 1.6 | 76.9 | 126.8 | 16.4 | 47.2 | 774.7 |
| CA2 rad | 5.2 | 12.1 | 63.1 | 7.7 | 0.0 | 0.0 | 2.9 | 0.0 | 0.0 | 2.5 | 0.0 | 0.0 | 1.5 | 55.5 | 84.1 | 19.9 | 7.4 | 147.2 |
| CA2 Im | 3.4 | 0.0 | 0.0 | 3.9 | 0.0 | 0.0 | 1.5 | 0.0 | 0.0 | 2.3 | 0.0 | 0.0 | 2.0 | 0.0 | 0.0 | 13.2 | 0.0 | 0.0 |
| CA3 ori | 7.1 | 154.4 | 1102.6 | 14.9 | 25.2 | 376.4 | 14.1 | 155.2 | 2193.5 | 8.9 | 161.4 | 1435.1 | 10.7 | 101.1 | 1080.3 | 55.8 | 110.9 | 6187.9 |
| CA3 pyr | 27.3 | 214.6 | 5851.4 | 77.7 | 58.3 | 4532.5 | 32.5 | 116.5 | 3787.3 | 28.7 | 234.5 | 6726.4 | 25.0 | 344.0 | 8607.6 | 191.2 | 154.3 | 29505.2 |
| CA3 rad | 10.3 | 112.4 | 1156.2 | 9.9 | 14.5 | 142.6 | 8.0 | 33.2 | 266.2 | 3.7 | 54.7 | 201.0 | 3.4 | 0.0 | 0.0 | 35.2 | 50.2 | 1766.0 |
| CA3 Im | 2.4 | 58.5 | 138.5 | 4.4 | 20.3 | 88.6 | 1.9 | 0.0 | 0.0 | 1.2 | 0.0 | 0.0 | 1.7 | 0.0 | 0.0 | 11.6 | 19.6 | 227.1 |
| DG mol | 30.4 | 2.0 | 59.5 | 41.0 | 2.0 | 83.9 | 32.8 | 0.0 | 0.0 | 26.0 | 1.7 | 45.5 | 27.9 | 0.0 | 0.0 | 158.2 | 1.2 | 188.9 |
| DG gra | 9.5 | 30.1 | 285.3 | 44.6 | 9.2 | 412.3 | 10.1 | 0.0 | 0.0 | 11.4 | 21.5 | 245.9 | 7.5 | 0.0 | 0.0 | 83.1 | 11.4 | 943.5 |
| DG hil | 10.7 | 183.9 | 1968.4 | 15.1 | 64.1 | 969.0 | 9.6 | 240.3 | 2311.6 | 12.1 | 163.1 | 1977.0 | 8.1 | 170.5 | 1377.5 | 55.7 | 154.6 | 8603.5 |
| Sum of all |  |  |  |  |  |  |  |  |  |  |  |  |  |  |  | 1106.5 |  | 71350.1 |

Subject: SKO27, Immunohistochemistry: CR
| Block | 1 from 0 um - (Head) |  |  | 2 from 3980 um - |  |  | 3 from 12580 um - |  |  | 4 from 22180 um - |  |  | 5 from 31960 um - |  |  | Total |  |  |
| --- | --- | --- | --- | --- | --- | --- | --- | --- | --- | --- | --- | --- | --- | --- | --- | --- | --- | --- |
| SKO27 CR | Volume [mm<sup>3</sup>] | Density [cell/mm<sup>3</sup>] | Abs. No | Volume [mm<sup>3</sup>] | Density [cell/mm<sup>3</sup>] | Abs. No | Volume [mm<sup>3</sup>] | Density [cell/mm<sup>3</sup>] | Abs. No | Volume [mm<sup>3</sup>] | Density [cell/mm<sup>3</sup>] | Abs. No | Volume [mm<sup>3</sup>] | Density [cell/mm<sup>3</sup>] | Abs. No | Volume [mm<sup>3</sup>] | Density [cell/mm<sup>3</sup>] | Abs. No |
| CA1 ori | 28.7 | 177.1 | 5086.9 | 13.2 | 89.5 | 1177.9 | 10.4 | 47.6 | 495.8 | 13.0 | 97.2 | 1264.0 | 9.8 | 115.7 | 1129.1 | 75.1 | 121.9 | 9153.7 |
| CA1pyr | 68.9 | 128.9 | 8882.1 | 42.3 | 140.1 | 5933.0 | 19.0 | 94.3 | 1791.8 | 20.1 | 162.2 | 3253.1 | 27.5 | 250.6 | 6894.9 | 177.8 | 150.5 | 26754.9 |
| CA1 rad | 31.1 | 290.2 | 9038.0 | 17.1 | 337.0 | 5758.3 | 19.6 | 326.8 | 6417.7 | 15.4 | 592.2 | 9109.5 | 14.8 | 1357.7 | 20083.1 | 98.0 | 514.1 | 50406.7 |
| CA1 Im | 53.3 | 773.7 | 41254.9 | 16.9 | 566.6 | 9588.9 | 7.3 | 391.4 | 2855.7 | 10.2 | 320.7 | 3278.0 | 11.2 | 389.0 | 4337.6 | 98.9 | 619.9 | 61315.2 |
| CA2 ori | 5.9 | 393.1 | 2319.6 | 4.2 | 191.3 | 804.3 | 2.0 | 50.0 | 100.8 | 3.5 | 83.5 | 296.4 | 0.9 | 85.9 | 76.1 | 16.6 | 217.3 | 3597.1 |
| CA2 pyr | 4.3 | 321.6 | 1372.6 | 5.3 | 42.6 | 224.4 | 2.3 | 42.0 | 96.6 | 2.9 | 23.7 | 69.6 | 1.6 | 20.0 | 33.0 | 16.4 | 109.4 | 1796.2 |
| CA2 rad | 5.2 | 161.9 | 842.5 | 7.7 | 86.7 | 664.7 | 2.9 | 110.6 | 323.1 | 2.5 | 45.1 | 115.0 | 1.5 | 133.8 | 202.7 | 19.9 | 108.2 | 2148.1 |
| CA2 Im | 3.4 | 361.9 | 1243.5 | 3.9 | 290.4 | 1141.9 | 1.5 | 99.7 | 150.4 | 2.3 | 0.0 | 0.0 | 2.0 | 114.1 | 223.6 | 13.2 | 209.3 | 2759.4 |
| CA3 ori | 7.1 | 120.4 | 859.5 | 14.9 | 190.7 | 2847.9 | 14.1 | 25.4 | 359.4 | 8.9 | 77.3 | 687.7 | 10.7 | 127.3 | 1361.0 | 55.8 | 109.6 | 6115.5 |
| CA3 pyr | 27.3 | 164.1 | 4474.2 | 77.7 | 48.7 | 3786.5 | 32.5 | 18.6 | 604.2 | 28.7 | 104.0 | 2983.3 | 25.0 | 202.8 | 5076.0 | 191.2 | 88.5 | 16924.1 |
| CA3 rad | 10.3 | 215.3 | 2215.4 | 9.9 | 134.4 | 1325.0 | 8.0 | 50.2 | 402.2 | 3.7 | 85.3 | 313.6 | 3.4 | 93.6 | 314.2 | 35.2 | 129.9 | 4570.4 |
| CA3 Im | 2.4 | 286.9 | 679.3 | 4.4 | 167.6 | 732.1 | 1.9 | 0.0 | 0.0 | 1.2 | 0.0 | 0.0 | 1.7 | 174.2 | 294.7 | 11.6 | 147.5 | 1706.1 |
| DG mol | 30.4 | 125.8 | 3824.0 | 41.0 | 116.5 | 4779.6 | 32.8 | 94.5 | 3104.9 | 26.0 | 132.0 | 3436.7 | 27.9 | 215.8 | 6019.2 | 158.2 | 133.8 | 21164.4 |
| DG gra | 9.5 | 233.1 | 2211.7 | 44.6 | 313.1 | 13963.0 | 10.1 | 301.7 | 3039.6 | 11.4 | 473.0 | 5401.5 | 7.5 | 1017.4 | 7649.3 | 83.1 | 388.3 | 32265.1 |
| DG hil | 10.7 | 343.6 | 3677.6 | 15.1 | 397.1 | 6007.6 | 9.6 | 394.2 | 3791.0 | 12.1 | 606.3 | 7350.2 | 8.1 | 891.2 | 7199.7 | 55.7 | 503.6 | 28026.1 |
| Sum of all |  |  |  |  |  |  |  |  |  |  |  |  |  |  |  | 1106.5 |  | 268703.0 |

Subject: SKO31, Immunohistochemistry: PV
| Block | 1 from 0 um - (Head) |  |  | 2 from 9560 um - |  |  | 3 from 16560 um - |  |  | 4 from 23860 um - |  |  | 5 from 32160 um - |  |  | Total |  |  |
| --- | --- | --- | --- | --- | --- | --- | --- | --- | --- | --- | --- | --- | --- | --- | --- | --- | --- | --- |
| SKO31 PV | Volume [mm<sup>3</sup>] | Density [cell/mm<sup>3</sup>] | Abs. No | Volume [mm<sup>3</sup>] | Density [cell/mm<sup>3</sup>] | Abs. No | Volume [mm<sup>3</sup>] | Density [cell/mm<sup>3</sup>] | Abs. No | Volume [mm<sup>3</sup>] | Density [cell/mm<sup>3</sup>] | Abs. No | Volume [mm<sup>3</sup>] | Density [cell/mm<sup>3</sup>] | Abs. No | Volume [mm<sup>3</sup>] | Density [cell/mm<sup>3</sup>] | Abs. No |
| CA1 ori | 50.7 | 82.8 | 4196.0 | 27.3 | 97.5 | 2656.7 | 20.3 | 91.3 | 1851.2 | 11.8 | 31.7 | 374.8 | 11.7 | 201.9 | 2362.5 | 121.7 | 94.0 | 11441.3 |
| CA1 pyr | 117.6 | 30.1 | 3539.2 | 50.8 | 43.9 | 2229.9 | 43.1 | 47.8 | 2062.7 | 28.8 | 99.7 | 2876.7 | 33.0 | 167.5 | 5531.3 | 273.4 | 59.4 | 16239.7 |
| CA1 rad | 45.1 | 3.5 | 158.1 | 27.3 | 0.0 | 0.0 | 17.7 | 3.2 | 57.2 | 9.3 | 8.1 | 75.5 | 15.9 | 13.8 | 218.3 | 115.2 | 4.4 | 509.1 |
| CA1 Im | 72.3 | 8.7 | 632.0 | 56.4 | 26.1 | 1470.9 | 23.0 | 7.7 | 176.2 | 9.3 | 8.0 | 74.0 | 17.7 | 31.0 | 549.5 | 178.7 | 16.2 | 2902.7 |
| CA2 ori | 11.2 | 39.1 | 439.5 | 4.6 | 118.4 | 544.8 | 2.8 | 73.5 | 203.1 | 3.1 | 24.1 | 75.2 | 3.2 | 221.1 | 709.1 | 24.9 | 79.1 | 1971.6 |
| CA2 pyr | 45.8 | 22.0 | 1007.4 | 8.9 | 59.3 | 526.9 | 3.9 | 99.7 | 384.9 | 5.3 | 136.1 | 726.4 | 5.9 | 195.6 | 1160.8 | 69.8 | 54.5 | 3806.5 |
| CA2 rad | 20.3 | 0.0 | 0.0 | 9.5 | 9.5 | 90.4 | 3.9 | 0.0 | 0.0 | 3.2 | 0.0 | 0.0 | 3.5 | 0.0 | 0.0 | 40.4 | 2.2 | 90.4 |
| CA2 Im | 30.7 | 9.3 | 284.5 | 7.9 | 23.9 | 188.5 | 2.1 | 0.0 | 0.0 | 3.7 | 0.0 | 0.0 | 2.4 | 0.0 | 0.0 | 46.8 | 10.1 | 473.0 |
| CA3 ori | 8.9 | 4.7 | 42.1 | 19.3 | 29.1 | 562.2 | 13.2 | 21.9 | 288.0 | 9.4 | 18.0 | 169.4 | 8.9 | 132.1 | 1181.1 | 59.7 | 37.5 | 2242.7 |
| CA3 pyr | 47.8 | 8.8 | 421.0 | 62.3 | 31.3 | 1949.8 | 39.2 | 23.7 | 927.3 | 38.7 | 25.0 | 967.0 | 38.4 | 48.5 | 1861.2 | 226.4 | 27.1 | 6126.2 |
| CA3 rad | 0.0 | 0.0 | 0.0 | 11.6 | 7.2 | 83.9 | 4.5 | 10.6 | 48.0 | 3.0 | 22.1 | 67.3 | 5.0 | 0.0 | 0.0 | 24.2 | 8.2 | 199.1 |
| CA3 Im | 0.0 | 0.0 | 0.0 | 7.4 | 5.9 | 43.6 | 0.7 | 0.0 | 0.0 | 2.4 | 17.0 | 40.3 | 1.5 | 0.0 | 0.0 | 12.0 | 7.0 | 83.9 |
| DG mol | 28.0 | 3.0 | 84.2 | 35.2 | 7.9 | 279.5 | 36.3 | 5.9 | 215.2 | 35.7 | 0.0 | 0.0 | 34.0 | 0.0 | 0.0 | 169.1 | 3.4 | 578.9 |
| DG gra | 9.7 | 4.3 | 42.1 | 12.1 | 5.5 | 66.1 | 9.6 | 11.5 | 110.6 | 9.0 | 40.9 | 366.2 | 10.5 | 80.5 | 845.7 | 50.9 | 28.1 | 1430.8 |
| DG hil | 12.5 | 13.4 | 168.4 | 15.8 | 33.1 | 524.6 | 11.3 | 19.6 | 220.3 | 8.1 | 51.5 | 417.8 | 9.2 | 83.8 | 769.1 | 56.9 | 36.9 | 2100.2 |
| Sum of all |  |  |  |  |  |  |  |  |  |  |  |  |  |  |  | 1470.3 |  | 50196.2 |

Subject: SKO31, Immunohistochemistry: SOM
| Block | 1 from 0 um - (Head) |  |  | 2 from 9560 um - |  |  | 3 from 16560 um - |  |  | 4 from 23860 um - |  |  | 5 from 32160 um - |  |  | Total |  |  |
| --- | --- | --- | --- | --- | --- | --- | --- | --- | --- | --- | --- | --- | --- | --- | --- | --- | --- | --- |
| SKO31 SOM | Volume [mm<sup>3</sup>] | Density [cell/mm<sup>3</sup>] | Abs. No | Volume [mm<sup>3</sup>] | Density [cell/mm<sup>3</sup>] | Abs. No | Volume [mm<sup>3</sup>] | Density [cell/mm<sup>3</sup>] | Abs. No | Volume [mm<sup>3</sup>] | Density [cell/mm<sup>3</sup>] | Abs. No | Volume [mm<sup>3</sup>] | Density [cell/mm<sup>3</sup>] | Abs. No | Volume [mm<sup>3</sup>] | Density [cell/mm<sup>3</sup>] | Abs. No |
| CA1 ori | 50.7 | 102.0 | 5171.4 | 27.3 | 53.2 | 1449.6 | 20.3 | 259.5 | 5260.9 | 11.8 | 75.7 | 895.3 | 11.7 | 135.6 | 1586.0 | 121.7 | 118.0 | 14363.1 |
| CA1 pyr | 117.6 | 7.2 | 852.6 | 50.8 | 7.6 | 384.4 | 43.1 | 19.9 | 860.2 | 28.8 | 23.2 | 668.4 | 33.0 | 16.7 | 552.4 | 273.4 | 12.1 | 3318.0 |
| CA1 rad | 45.1 | 4.1 | 185.0 | 27.3 | 4.4 | 119.6 | 17.7 | 11.2 | 197.5 | 9.3 | 0.0 | 0.0 | 15.9 | 7.7 | 122.0 | 115.2 | 5.4 | 624.1 |
| CA1 Im | 72.3 | 2.9 | 207.0 | 56.4 | 0.0 | 0.0 | 23.0 | 0.0 | 0.0 | 9.3 | 0.0 | 0.0 | 17.7 | 0.0 | 0.0 | 178.7 | 1.2 | 207.0 |
| CA2 ori | 11.2 | 126.3 | 1419.8 | 4.6 | 182.2 | 838.2 | 2.8 | 124.1 | 342.6 | 3.1 | 53.7 | 167.3 | 3.2 | 183.1 | 587.3 | 24.9 | 134.6 | 3355.3 |
| CA2 pyr | 45.8 | 23.1 | 1058.0 | 8.9 | 13.1 | 116.6 | 3.9 | 13.4 | 51.7 | 5.3 | 15.3 | 81.6 | 5.9 | 45.2 | 268.3 | 69.8 | 22.6 | 1576.3 |
| CA2 rad | 20.3 | 10.2 | 207.7 | 9.5 | 14.0 | 133.0 | 3.9 | 0.0 | 0.0 | 3.2 | 0.0 | 0.0 | 3.5 | 0.0 | 0.0 | 40.4 | 8.4 | 340.7 |
| CA2 Im | 30.7 | 5.2 | 160.1 | 7.9 | 0.0 | 0.0 | 2.1 | 0.0 | 0.0 | 3.7 | 0.0 | 0.0 | 2.4 | 0.0 | 0.0 | 46.8 | 3.4 | 160.1 |
| CA3 ori | 8.9 | 47.7 | 425.5 | 19.3 | 119.9 | 2318.4 | 13.2 | 185.4 | 2439.4 | 9.4 | 75.6 | 710.6 | 8.9 | 161.3 | 1441.7 | 59.7 | 122.8 | 7335.6 |
| CA3 pyr | 47.8 | 162.6 | 7776.2 | 62.3 | 104.6 | 6508.7 | 39.2 | 104.9 | 4106.2 | 38.7 | 80.6 | 3118.9 | 38.4 | 207.6 | 7974.2 | 226.4 | 130.3 | 29484.2 |
| CA3 rad | 0.0 | 0.0 | 0.0 | 11.6 | 0.0 | 0.0 | 4.5 | 33.4 | 151.3 | 3.0 | 0.0 | 0.0 | 5.0 | 28.0 | 139.2 | 24.2 | 12.0 | 290.4 |
| CA3 Im | 0.0 | 0.0 | 0.0 | 7.4 | 0.0 | 0.0 | 0.7 | 15.7 | 11.1 | 2.4 | 0.0 | 0.0 | 1.5 | 0.0 | 0.0 | 12.0 | 0.9 | 11.1 |
| DG mol | 28.0 | 0.0 | 0.0 | 35.2 | 4.5 | 156.8 | 36.3 | 2.2 | 79.7 | 35.7 | 0.0 | 0.0 | 34.0 | 0.0 | 0.0 | 169.1 | 1.4 | 236.5 |
| DG gra | 9.7 | 0.0 | 0.0 | 12.1 | 6.1 | 73.6 | 9.6 | 24.7 | 238.1 | 9.0 | 0.0 | 0.0 | 10.5 | 7.9 | 82.8 | 50.9 | 7.7 | 394.5 |
| DG hil | 12.5 | 128.5 | 1610.8 | 15.8 | 159.4 | 2524.1 | 11.3 | 140.9 | 1585.3 | 8.1 | 51.3 | 416.0 | 9.2 | 155.9 | 1430.6 | 56.9 | 133.0 | 7566.8 |
| Sum of all |  |  |  |  |  |  |  |  |  |  |  |  |  |  |  | 1470.3 |  | 69263.7 |

Subject: SKO31, Immunohistochemistry: CR
| Block | 1 from 0 um - (Head) |  |  | 2 from 9560 um - |  |  | 3 from 16560 um - |  |  | 4 from 23860 um - |  |  | 5 from 32160 um - |  |  | Total |  |  |
| --- | --- | --- | --- | --- | --- | --- | --- | --- | --- | --- | --- | --- | --- | --- | --- | --- | --- | --- |
| SKO31 CR | Volume [mm<sup>3</sup>] | Density [cell/mm<sup>3</sup>] | Abs. No | Volume [mm<sup>3</sup>] | Density [cell/mm<sup>3</sup>] | Abs. No | Volume [mm<sup>3</sup>] | Density [cell/mm<sup>3</sup>] | Abs. No | Volume [mm<sup>3</sup>] | Density [cell/mm<sup>3</sup>] | Abs. No | Volume [mm<sup>3</sup>] | Density [cell/mm<sup>3</sup>] | Abs. No | Volume [mm<sup>3</sup>] | Density [cell/mm<sup>3</sup>] | Abs. No |
| CA1 ori | 50.7 | 93.1 | 4717.3 | 27.3 | 30.2 | 823.6 | 20.3 | 61.0 | 1237.6 | 11.8 | 27.2 | 322.0 | 11.7 | 134.4 | 1572.5 | 121.7 | 71.2 | 8672.9 |
| CA1 pyr | 117.6 | 103.3 | 12147.2 | 50.8 | 99.2 | 5035.5 | 43.1 | 101.6 | 4379.7 | 28.8 | 84.9 | 2450.4 | 33.0 | 134.9 | 4453.4 | 273.4 | 104.1 | 28466.2 |
| CA1 rad | 45.1 | 276.3 | 12449.9 | 27.3 | 200.4 | 5463.5 | 17.7 | 233.5 | 4129.5 | 9.3 | 484.5 | 4526.3 | 15.9 | 872.4 | 13836.0 | 115.2 | 350.7 | 40405.2 |
| CA1 Im | 72.3 | 822.8 | 59489.6 | 56.4 | 621.5 | 35077.4 | 23.0 | 637.7 | 14657.9 | 9.3 | 442.4 | 4106.9 | 17.7 | 521.3 | 9239.6 | 178.7 | 685.8 | 122571.4 |
| CA2 ori | 11.2 | 73.4 | 825.4 | 4.6 | 78.6 | 361.6 | 2.8 | 193.8 | 535.1 | 3.1 | 132.0 | 411.5 | 3.2 | 138.1 | 442.9 | 24.9 | 103.4 | 2576.5 |
| CA2 pyr | 45.8 | 126.8 | 5809.3 | 8.9 | 69.3 | 615.6 | 3.9 | 55.7 | 215.0 | 5.3 | 109.8 | 585.7 | 5.9 | 22.5 | 133.4 | 69.8 | 105.4 | 7359.0 |
| CA2 rad | 20.3 | 388.5 | 7901.6 | 9.5 | 109.8 | 1043.8 | 3.9 | 54.3 | 210.6 | 3.2 | 63.3 | 200.7 | 3.5 | 183.8 | 648.1 | 40.4 | 247.5 | 10004.9 |
| CA2 Im | 30.7 | 779.1 | 23940.0 | 7.9 | 250.0 | 1975.7 | 2.1 | 146.8 | 308.3 | 3.7 | 100.9 | 370.4 | 2.4 | 53.0 | 129.5 | 46.8 | 570.5 | 26724.0 |
| CA3 ori | 8.9 | 0.0 | 0.0 | 19.3 | 110.9 | 2143.1 | 13.2 | 318.5 | 4191.8 | 9.4 | 128.0 | 1202.8 | 8.9 | 253.1 | 2262.5 | 59.7 | 164.0 | 9800.3 |
| CA3 pyr | 47.8 | 68.2 | 3259.2 | 62.3 | 76.3 | 4746.9 | 39.2 | 97.3 | 3808.6 | 38.7 | 42.8 | 1655.9 | 38.4 | 107.6 | 4134.2 | 226.4 | 77.8 | 17604.9 |
| CA3 rad | 0.0 | 0.0 | 0.0 | 11.6 | 144.4 | 1681.2 | 4.5 | 145.6 | 659.0 | 3.0 | 326.0 | 992.6 | 5.0 | 205.0 | 1017.1 | 24.2 | 179.9 | 4349.7 |
| CA3 Im | 0.0 | 0.0 | 0.0 | 7.4 | 227.8 | 1695.9 | 0.7 | 154.7 | 109.6 | 2.4 | 129.0 | 306.7 | 1.5 | 142.4 | 215.2 | 12.0 | 193.3 | 2327.3 |
| DG mol | 28.0 | 120.6 | 3373.2 | 35.2 | 81.9 | 2883.3 | 36.3 | 135.5 | 4916.5 | 35.7 | 187.2 | 6675.5 | 34.0 | 204.2 | 6936.6 | 169.1 | 146.6 | 24785.1 |
| DG gra | 9.7 | 603.7 | 5866.0 | 12.1 | 585.9 | 7085.7 | 9.6 | 559.2 | 5395.7 | 9.0 | 624.4 | 5591.5 | 10.5 | 925.3 | 9725.4 | 50.9 | 661.0 | 33664.3 |
| DG hil | 12.5 | 414.5 | 5196.1 | 15.8 | 521.1 | 8252.0 | 11.3 | 666.0 | 7494.1 | 8.1 | 420.6 | 3412.0 | 9.2 | 820.4 | 7526.8 | 56.9 | 560.2 | 31881.1 |
| Sum of all |  |  |  |  |  |  |  |  |  |  |  |  |  |  |  | 1470.3 |  | 371192.7 |

**Extended Data Table 2:** Absolute number of interneurons in the anterior, medial and posterior third of the hippocampus in subjects SKO24, 25, 27 and 31.

**Absolute number of interneurons in the different parts of the hippocampus**
| Subject | Anterior |  |  | Medial |  |  | Posterior |  |  |
| --- | --- | --- | --- | --- | --- | --- | --- | --- | --- |
|  | PV | SOM | CR | PV | SOM | CR | PV | SOM | CR |
| **SKO24** | 10878.1 | 48314.7 | 51741.3 | 13488.9 | 50456.3 | 68909.2 | 24283.9 | 53823.1 | 85054.6 |
| **SKO25** | 9026.7 | 36989.3 | 63982.8 | 12743.6 | 44041.0 | 76515.5 | 18468.9 | 49430.9 | 91949.7 |
| **SKO27** | 11276.1 | 28168.4 | 100956.2 | 13470.9 | 13591.3 | 62096.7 | 33986.2 | 29590.4 | 105650.1 |
| **SKO31** | 10780.6 | 18669.3 | 141897.0 | 14931.8 | 23176.4 | 109745.5 | 24483.8 | 27418.1 | 119550.2 |

**Extended Data table 3:** Density of interneurons [cell/mm^3^] in the anterior, medial and posterior third of the hippocampus in subjects SKO24, 25, 27 and 31.

**Density of interneurons [cell/mm<sup>3</sup>] in the different parts of the hippocampus**
| Subject | Anterior |  |  | Medial |  |  | Posterior |  |  |
| --- | --- | --- | --- | --- | --- | --- | --- | --- | --- |
|  | PV | SOM | CR | PV | SOM | CR | PV | SOM | CR |
| **SKO24** | 58.0 | 257.5 | 275.8 | 71.9 | 268.9 | 367.3 | 129.4 | 286.9 | 453.3 |
| **SKO25** | 37.7 | 154.7 | 267.6 | 53.3 | 184.2 | 320.0 | 77.2 | 206.7 | 384.5 |
| **SKO27** | 30.6 | 76.4 | 273.7 | 36.5 | 36.8 | 168.4 | 92.1 | 80.2 | 286.4 |
| **SKO31** | 22.0 | 38.1 | 289.5 | 30.5 | 47.3 | 223.9 | 50.0 | 55.9 | 243.9 |

